# Appraising causal relationships of dietary, nutritional and physical-activity exposures with overall and aggressive prostate cancer: two-sample Mendelian randomization study based on 79,148 prostate cancer cases and 61,106 controls

**DOI:** 10.1101/674820

**Authors:** Nabila Kazmi, Philip Haycock, Konstantinos Tsilidis, Brigid M. Lynch, Therese Truong, the PRACTICAL consortium, CRUK, BPC3, CAPS, PEGASUS, Richard M Martin, Sarah Lewis

## Abstract

**Background:** Prostate cancer is the second most common male cancer worldwide, but there is substantial geographical variation suggesting a potential role for modifiable risk factors in prostate carcinogenesis.

**Methods:** We identified previously reported prostate cancer risk factors from the World Cancer Research Fund’s (WCRF) systematic appraisal of the global evidence (2018). We assessed whether each identified risk factor was causally associated with risk of overall (79,148 cases and 61,106 controls) or aggressive (15,167 cases and 58,308 controls) prostate cancer using Mendelian randomization (MR) based on genome wide association study (GWAS) summary statistics from the PRACTICAL and GAME-ON/ELLIPSE consortia. We assessed evidence for replication in UK Biobank (7,844 prostate cancer cases and 204,001 controls).

**Findings:** WCRF identified 57 potential risk factors, of which 22 could be instrumented for MR analyses using single nucleotide polymorphisms (SNPs). In MR analyses for overall prostate cancer, we identified evidence compatible with causality for the following risk factors (odds ratio [OR] per standard deviation increase; 95% confidence interval): accelerometer-measured physical-activity, OR=0.49 (0.33-0.72; p=0.0003); serum iron, OR=0.92 (0.86-0.98; p=0.007); body mass index (BMI), OR=0.90 (0.84-0.97; p=0.003); and mono-unsaturated fat, OR=1.11 (1.02-1.20; p=0.02). Findings in our replication analyses in UK Biobank were compatible with our main analyses (albeit with wide confidence intervals). In MR analysis, height was positively associated with aggressive prostate cancer risk: OR=1.07 (1.01-1.15; p=0.03).

**Interpretation:** The results for physical-activity, serum iron, BMI, mono-unsaturated fat and height are compatible with causality for prostate cancer but more research is needed to rule out violations of MR assumptions for some risk factors. The results suggest that interventions aimed at increasing physical activity may reduce prostate cancer risk, but the direction of effects of BMI, and iron are at odds with their effects on other diseases, so the overall public health impact of intervening on these need to be considered.

**Funding:** World Cancer Research Fund International (2015/1421), Cancer Research UK program grant (C18281/A19169), National Institute for Health Research, Bristol Biomedical Research Centre, and Victorian Cancer Agency (MCRF18005).

## Research in context

### Evidence before this study

The World Cancer Research Fund (WCRF) Continuous Update Project (CUP) has reported associations between diet, nutrition and physical activity, and prostate cancer using observational studies. Establishing causality is an important step in the development of prevention strategies but is challenging because inference from observational studies is limited by often intractable biases, including measurement error, reverse causation and residual, or unmeasured confounding.

### Added value of this study

Mendelian randomization (MR) is a methodological approach to addressing measurement error, reverse causation and confounding within observational studies, based on ‘instrumental variable’ (IV) analysis. We systematically applied two-sample MR to appraise the evidence for a causal link between previously reported lifestyle and anthropometric risk factors (from WCRF report) with overall prostate cancer risk and aggressive prostate cancer risk.

### Implications of all the available evidence

Our MR analyses showed that physical activity, BMI, and serum iron levels were inversely associated with overall prostate cancer risk and mono-unsaturated fat levels were positively associated and that these effects were likely to be causal. For these risk factors, the direction of association was consistent for aggressive prostate cancer risk. In addition, our MR analyses showed that height was positively associated with aggressive prostate cancer risk.

## Introduction

In 2012, 1.1 million men were diagnosed with prostate cancer, making it the second most common male cancer worldwide^1,2^. There is wide global variation in prostate cancer incidence, with almost 70% of cases occurring in more developed regions of the world^2^. This variation is thought in part to be related to the intensity of prostate-specific antigen (PSA) based screening practices^3,4^, although migration studies suggest an influence of environmental and lifestyle factors^5,6^. Established risk factors include advanced age, ethnicity and family history of prostate cancer^7,8^. In addition, several lifestyle and anthropometric factors have been hypothesised to play an aetiological role, and measures of adiposity a prognostic role^8^. However, the epidemiological evidence to support a causal role for these potentially modifiable factors is weak. This is because inference from observational studies is limited by residual or unmeasured confounding, and other biases such as reverse causation and detection bias^9,10^.

Mendelian randomization (MR) is a methodological approach to addressing reverse causation and confounding within observational studies, based on long-established ‘instrumental variable’ (IV) principles^11^. MR exploits the random assortment of alleles through meiotic cell division at conception, and can be thought of as a natural experiment that generates conditions equivalent to randomised controlled trials, where randomised treatment arms are analogous to randomly assigned genetic subgroups^12–15^. At the population level, individuals defined by specific genotypes should on average only differ with respect to that genotype and its phenotypic consequences if certain IV assumptions are met. These assumptions are that the instrument is: i) robustly associated with the exposure it is acting as a proxy for; ii) independent of confounders; and iii) independent of the outcome conditional on the exposure (i.e. ‘no pleiotropy’ where a single locus influences the outcome through biological pathways that are independent of the exposure of interest)^16^. If these assumptions can be shown to have been met, then the genetic polymorphism can be used in an IV framework (i.e. MR) to provide an unconfounded and unbiased estimate of the causal association between the potentially modifiable risk factor and outcome of interest^17^. An extension of this methodology - two-sample MR - derives estimates for the required genotype-exposure and genotype-outcome associations from separate and non-overlapping samples of the same representative population^13,18^. Two-sample approaches exploit the rapidly growing availability of summary data from large consortia of genome-wide association studies (GWASs) and thus allow for greater sample sizes to improve statistical power^19^.

The aim of this study was to systematically apply two-sample MR analyses to appraise the evidence of a causal link between previously reported lifestyle and anthropometric risk factors with overall and aggressive prostate cancer.

## Methods

### Selection of risk factors

The World Cancer Research Fund (WCRF) Continuous Update Project (CUP) is a rigorous and systematic synthesis of the global scientific literature on diet, weight and physical-activity in relation to prostate cancer risk, based largely on observational epidemiological studies. We selected all potential risk factors for prostate cancer identified by the WCRF CUP reported in 2018^8^.

### Defining genetic instruments

We searched for each of the risk factors included in the WCRF report, using exact wording and synonyms, in both the GWAS catalog (www.ebi.ac.uk/gwas) and the MR-Base repository of full GWAS association statistics (www.mrbase.org)^20^. This was done to identify any studies that reported associations between single nucleotide polymorphisms (SNPs) and the specific risk factor of interest^20^. Further details of how we defined the genetic instruments and their properties are provided in the supplemental material, including; selection criteria for the SNP(s) to proxy each risk factor, how we standardised the beta coefficient and standard error (SE) for each SNP-exposure association, and how we calculated the proportion of variance (R^2^) in the risk factor explained by the SNP(s), the strength of the instrument represented by the F-statistic, and the power to detect an odds ratio (OR) of 1.2 (or conversely a protective OR of 0.80).

### Outcome trait

GWAS results for prostate cancer were obtained from fixed-effects meta-analyses based on individuals of European ancestry in the PRACTICAL and GAME-ON/ELLIPSE consortia (PRACTICAL: Prostate Cancer Association Group to Investigate Cancer-Associated Alterations in the Genome; GAME-ON: Genetic Associations and Mechanisms in Oncology; ELLIPSE: Elucidating Loci Involved in Prostate Cancer Susceptibility). The summary data were derived from a GWAS of overall prostate cancer on 79,148 cases and 61,106 controls^21^, and a GWAS of aggressive prostate cancer involving 15,167 cases and 58,308 controls^21^. Aggressive prostate cancer was defined as Gleason score ≥8, PSA >100 ng/mL, metastatic disease (M1), or death from prostate cancer.

### Two-sample MR analysis

We used the inverse-variance weighted (IVW) method as our primary MR analytical approach. The IVW method estimates the effect of the exposure on the outcome from the slope of the relationship between β_XG_ (SNP-exposure association) and β_YG_ (SNP-outcome association). This approach performs an inverse variance weighted meta-analysis of each Wald ratio^22^, effectively treating each SNP instrumenting a specific risk-factor as a valid natural experiment. We used a random effects IVW model by default, unless there was underdispersion in the causal estimates between SNPs, in which case a fixed effects model was used. The estimates from the random and fixed effects IVW models are the same but the variance for the random effects model is inflated to take into account heterogeneity between SNPs. For risk-factors that only had one SNP available as the instrument, we used the Wald ratio, which is equivalent to β_YG/_β_XG_ (where Y= outcome, G= gene and X=exposure).

In sensitivity analyses, we applied weighted median^23^, weighted mode^24^ and MR-Egger regression^25^ methods. The weighted median has the advantage that only half the SNPs need to be valid instruments (i.e. exhibiting no horizontal pleiotropy, no association with confounders, and a robust association with the exposure) for the causal effect estimate to be unbiased. The mode-based estimator clusters the SNPs into groups based on similarity of causal effects, and returns the causal effect estimate based on the cluster that has the largest number of SNPs. The weighted mode introduces an extra element similar to IVW and the weighted median, weighting each SNP’s contribution to the clustering by the inverse variance of its outcome effect.

The MR-Egger method is similar to the IVW approach but relaxes the ‘no horizontal pleiotropy’ assumption. MR-Egger regression allows a non-zero intercept in the relationship between multiple SNP-outcome and SNP-exposure associations, where the intercept provides a formal statistical test for the presence of directional (bias inducing) pleiotropy. The slope of the MR-Egger regression between multiple SNP-outcome and SNP-exposure associations can be considered as an unbiased causal effect between the risk factors and prostate cancer, assuming any horizontal pleiotropic effects are not correlated with the SNP-exposure effects (strength of the instrument). The MR pleiotropy residual sum and outlier test (MR-PRESSO) was also implemented to identify outlying genetic variants and analyses were re-run after excluding these variants^26^.Violations of the MR ‘no horizontal pleiotropy’ assumption were also assessed by visual inspection of funnel^27^, forest, scatter and leave-one-out plots, and tests of heterogeneity^28^ between SNPs making up a multi-allelic instrument^20^.

All analyses were conducted using the TwoSampleMR and MRInstruments R packages, curated by MR-Base, www.mrbase.org.

### Replication

The risk factors that showed suggestive evidence of association (p<0.05) with overall prostate cancer were assessed for replication among men in the UK Biobank prospective cohort of 7,844 prostate cancer cases and 204,001 controls, using two sample MR. The information on prostate cancer diagnosis was obtained from National Cancer Registries, UK (http://biobank.ndph.ox.ac.uk/showcase/field.cgi?id=40006), based on ICD10 code for prostate cancer C61. The GWAS results of this study are available on www.mrbase.org.

## Results

There were 57 potential risk factors for prostate cancer considered by the WCRF 2018 report^8^(Supplementary File 1; Table S1 and Table S2). The WCRF reported strong evidence that being obese (BMI, waist circumference and WHR) increases the risk of aggressive prostate cancer (Supplementary File 1; Table S2) and height increases the risk of prostate cancer (Supplementary File 1; Table S1 and S2). There was limited evidence that consumption of dairy products, diets high in calcium, and low plasma alpha-tocopherol and low plasm selenium concentrations increased prostate cancer risk. The evidence was too weak to draw any conclusions for the remaining risk factors. Of these 57 exposures, 22 could be analysed using MR because they had at least one SNP that was strongly associated with them (Table 1).

**Table 1.** Details of the instruments used to proxy risk factors for prostate cancer risk. BMI = body mass; WHR = waist-hip ratio; N is the sample size for the GWAS used to define the instruments; # SNPs represents the number of SNPs used within the instrument for each risk factor after clumping, harmonization and extraction of data from a GWAS of prostate cancer (# SNPs^1^ for overall prostate cancer risk and # SNPs^2^ for aggressive prostate cancer risk); Units and SD represent the analysis scale and standard deviation scale for betas and SE of SNPs for each risk factor(1 = the GWAS results were already on SD scale otherwise the SD of population mean), respectively; R^2^ represents the variance explained in the risk factor by the instrument; F indicates strength of the instrument used for each risk factor (a strong instrument is sometimes defined as an F-statistic >10); Power^2^ represents the power to detect an odds ratio of 1.2 for an association of the risk factor with overall prostate cancer at a significance level (P) of 0.05; Power^2^ represents the power to detect an odds ratio of 1.2 for an association of the risk factor with aggressive prostate cancer at a significance level (P) of 0.05.

| Risk factor | PubMed ID | N | # SNPs<sup>1</sup> | # SNPs<sup>2</sup> | Units | SD | R<sup>2</sup> | F | Power<sup>1</sup> | Power<sup>2</sup> |
| --- | --- | --- | --- | --- | --- | --- | --- | --- | --- | --- |
| *Anthropometrics and other measures* |  |  |  |  |  |  |  |  |  |  |
| Birth weight | 27680694 | 143677 | 46 | 45 | g | 1 | 1.69 | 53.54 | >99 | 53 |
| BMI | 30124842 | 681275 | 535 | 525 | kg/m<sup>2</sup> | 1 | 5.66 | 76.39 | >99 | 96 |
| Height | 25282103 | 253288 | 433 | 426 | m | 1 | 12.01 | 79.71 | >99 | >99 |
| Waist circumference | 25673412 | 232101 | 45 | 44 | cm | 1 | 0.95 | 49.60 | 94 | 33 |
| WHR | 25673412 | 212244 | 31 | 28 | ratio | 1 | 0.65 | 44.95 | 83 | 23 |
| *Circulating macro- and micro-nutrients* |  |  |  |  |  |  |  |  |  |  |
| Sugar/Sucrose (glucose) | 22885924 | 133010 | 15 | 15 | mmol/l | 0.73 | 0.52 | 46.65 | 74 | 21 |
| Monounsaturated fat | 27005778 | 13535 | 5 | 5 | mmol/l | 1 | 2.01 | 55.62 | >99 | 61 |
| Polyunsaturated fat | 27005778 | 13549 | 19 | 19 | mmol/l | 1 | 13.79 | 113.93 | >99 | >99 |
| Total fat | 27005778 | 13505 | 12 | 12 | mmol/l | 1 | 4.62 | 54.51 | >99 | 92 |
| Alpha-carotene | 28002826 | 433 | 3 | 3 | μmol/l | 0.23 | 22.08 | 40.53 | >99 | >99 |
| Beta-carotene | 19185284 | 3932 | 1 | 1 | μmol/l | 0.67 | 2.63 | 106.36 | >99 | 73 |
| Calcium | 24068962 | 61079 | 5 | 5 | mg/dl | 1 | 0.65 | 79.85 | 83 | 25 |
| Iron | 25352340 | 72958 | 5 | 5 | μmol/l | 1 | 2.20 | 328.85 | >99 | 66 |
| Lycopene | 26861389 | 441 | 1 | 1 | N/A | N/A | 8.34 | 39.93 | >99 | >99 |
| Phosphorous | 20558539 | 16264 | 4 | 4 | mg/dl | 0.49 | 1.52 | 62.82 | 99 | 50 |
| Retinol | 21878437 | 5006 | 2 | 2 | μg/l | 0.22 | 1.37 | 34.79 | 99 | 46 |
| Selenium | 25343990 | 9639 | 2 | 2 | μg/L | 0.18 | 1.44 | 70.33 | >99 | 48 |
| Vitamin D | 23393431 | 42024 | 4 | 4 | ng/ml | 10 | 2.35 | 253.15 | >99 | 68 |
| *Consumption of foods and drinks* |  |  |  |  |  |  |  |  |  |  |
| Alcohol | 30643251 | 941280 | 77 | 76 | drinks per week | 1 | 0.56 | 68.35 | 77 | 22 |
| Coffee | 25288136 | 91462 | 4 | 4 | cups/day | 1.96 | 0.45 | 103.79 | 68 | 19 |
| Dairy products (milk intake) | 29071499 | 74241 | 1 | 1 | glasses/week | 1.7 | 0.52 | 388.87 | 74 | 21 |
| *Physical-activity* |  |  |  |  |  |  |  |  |  |  |
| Average acceleration | 29899525 | 91084 | 2 | 2 | milli gravities (mg) | 8.14 | 0.10 | 44.68 | 20 | 7 |
| Sedentary behaviour | 30531941 | 91105 | 2 | 2 | proportion units | 1 | 0.08 | 34.91 | 17 | 6 |
| Fraction accelerations | 29899525 | 90667 | 1 | 1 | inverse-normalized fraction of accelerations | 1 | 0.04 | 39.06 | 11 | 5 |
| Sleep duration | 30531941 | 91105 | 7 | 7 | proportion units | 1 | 0.38 | 49.29 | 60 | 16 |

### Mendelian randomization results

Of the 22 potential risk factors examined in our study, only four showed evidence of an association with overall prostate cancer risk (Figure 1, Supplementary File 1; Table S3). Physical-activity, assessed as ‘average acceleration’ (OR per SD change: 0.49; 95% CI: 0.33, 0.72; P=0.0003), serum iron levels (OR: 0.92; 95% CI: 0.86, 0.98; P=0.007), and BMI (OR: 0.90; 95% CI: 0.84, 0.97; P=0.003) were inversely associated with overall prostate cancer risk. Circulating mono-unsaturated fat (OR: 1.11; 95% CI: 1.02, 1.20; P=0.02) was positively associated with overall prostate cancer risk. Compared to results from observational studies^8^ for overall prostate cancer risk (highest versus lowest total physical-activity; risk ratio (RR): 0.97; 95% CI: 0.9, 1.04), the estimate obtained by MR for the physical-activity measure (average acceleration) was much more strongly supportive of a protective effective (Figure 1). The WCRF reported strong evidence of association for increased body fatness (marked by BMI (RR: 1.08; 95% CI: 1.04, 1.12), waist circumference (RR: 1.12; 95% CI: 1.04, 1.21) and WHR (RR: 1.15; 95% CI: 1.03, 1.28)) with aggressive prostate cancer risk but these results were inconsistent with our MR results, which found evidence of protective association of BMI with overall prostate cancer risk. In the WCRF report (2018), the observational analysis reported no association between intake of mono-unsaturated fat and overall prostate cancer risk (RR: 1.00; 95% CI: 0.99, 1.01) but our MR-analysis found a positive association between circulating mono-unsaturated fat and overall prostate cancer risk.

**Figure 1.**
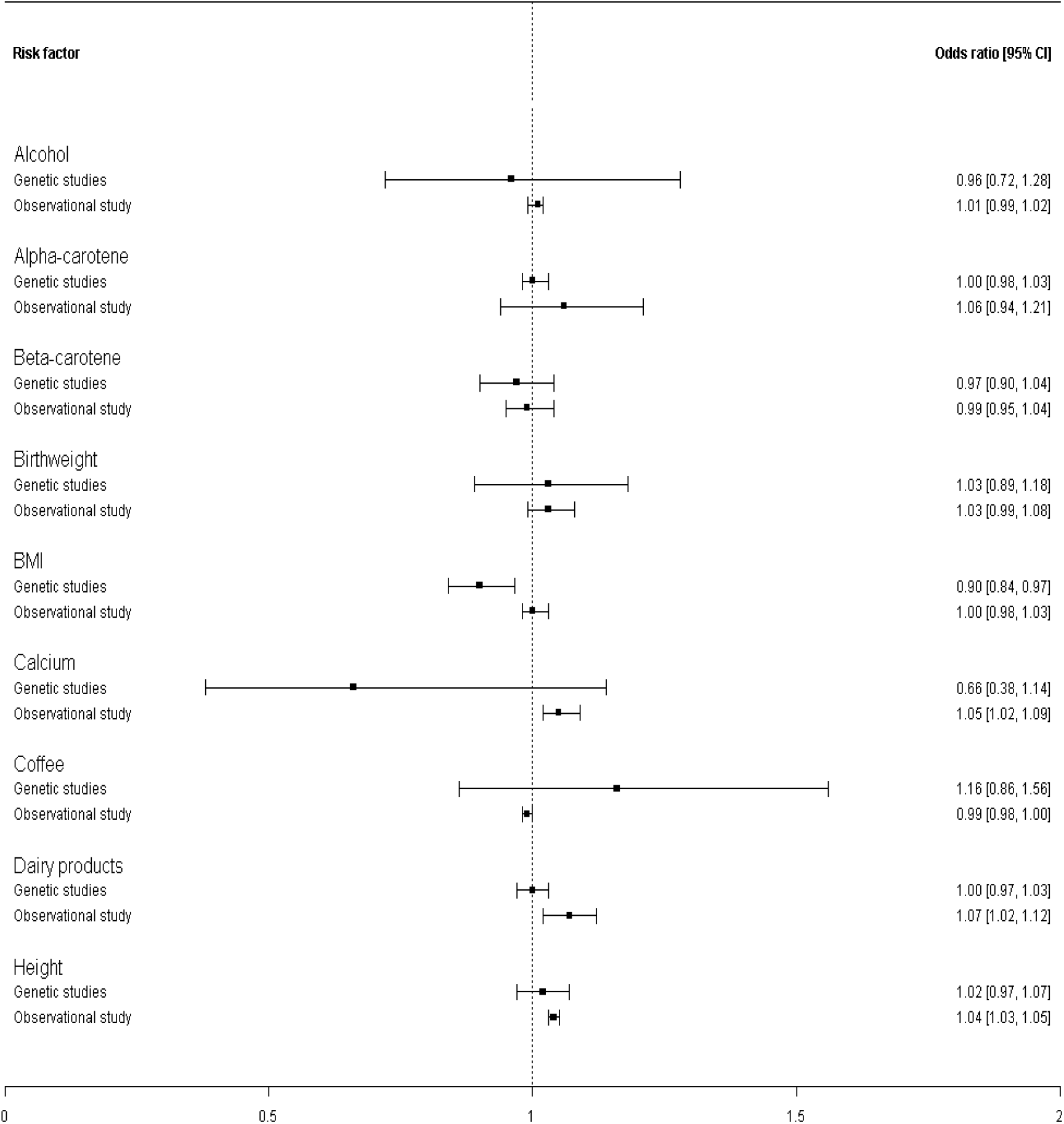
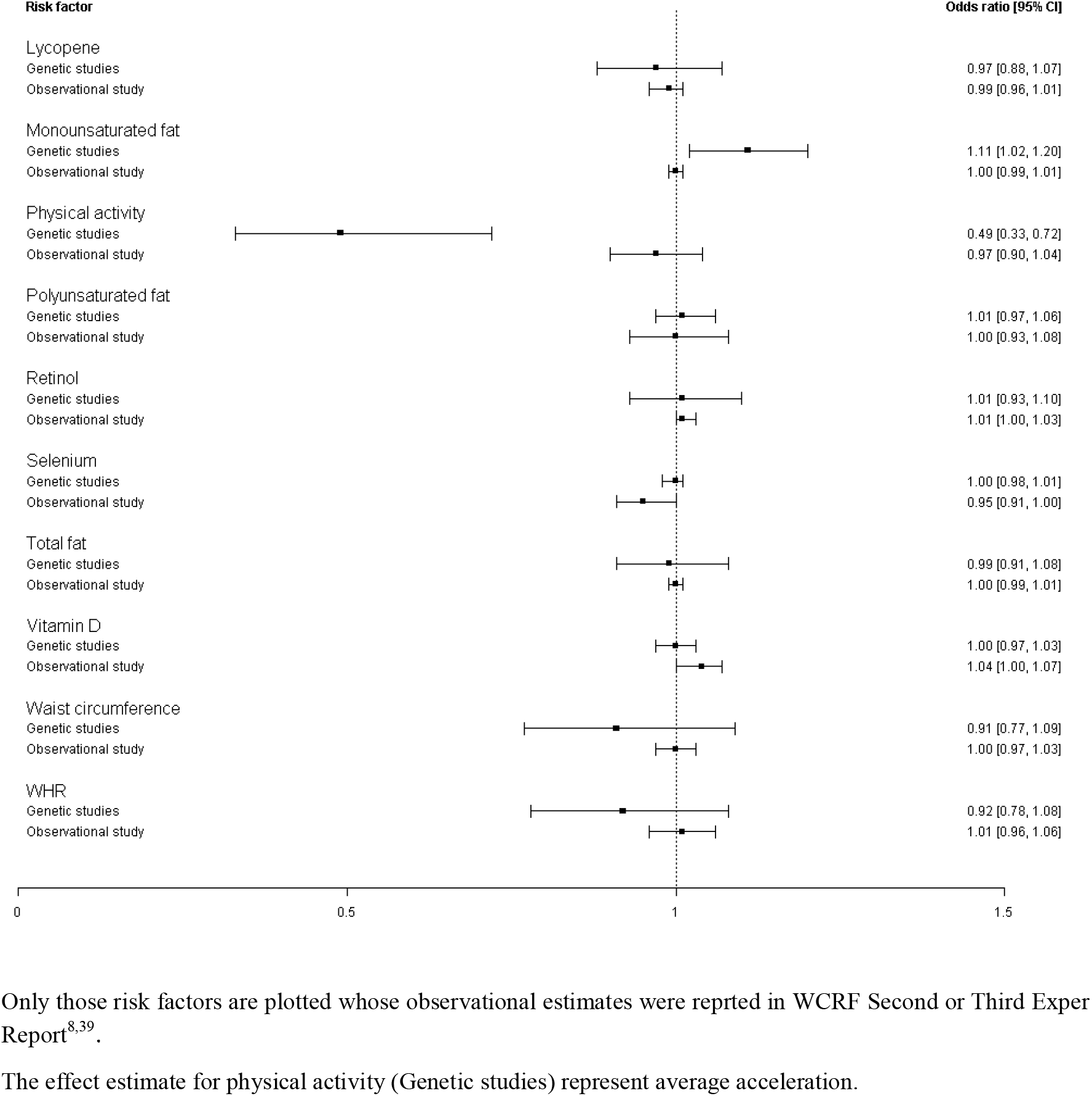
Comparison between observational and MR estimates for the risk factors and overall prostate cancer risk

None of the risk factors we examined showed strong evidence of association with aggressive prostate cancer although height showed weak evidence of increasing risk; OR: 1.07; 95% CI: 1.01, 1.15; P=0.03 (Figure 2, Supplementary File 1; Table S4).The observational studies also reported positive association of height with overall prostate cancer (OR: 1.04.; 95% CI: 1.03, 1.05; P=1.3×10^−15^) and aggressive prostate cancer^8^ (OR: 1.04; 95% CI: 1.02, 1.06; P=6.4×10^−5^). However, our MR analysis did not find evidence of association between height and overall prostate cancer risk. There was weak evidence for iron, and average acceleration, with effect-estimates being similar to those observed for overall prostate cancers. In the WCRF report, observational studies have reported positive effects of dairy products, calcium and low selenium concentration on overall prostate cancer, but we did not find strong evidence of associations with these in our MR analyses (Figure 1). However, the confidence intervals (CIs) for these risk factors were overlapping between observational and MR analyses. The power to detect the observationally reported effect size for these risk factors was >74%). Low alpha-tocopheral concentration was positively associated with prostate cancer risk in WCRF report but due to lack of an instrument it was not possible to conduct MR analyses.

**Figure 2.**
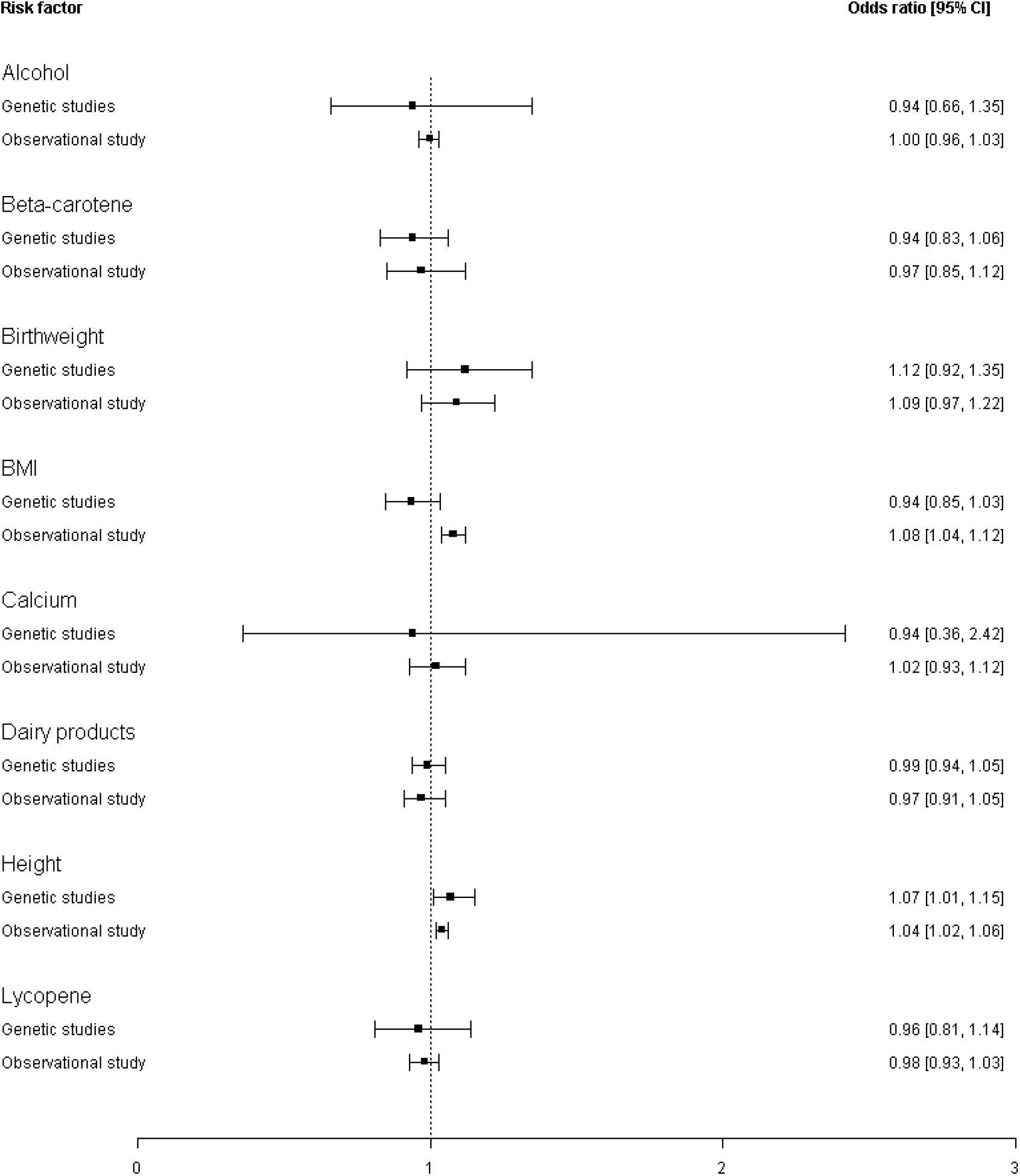
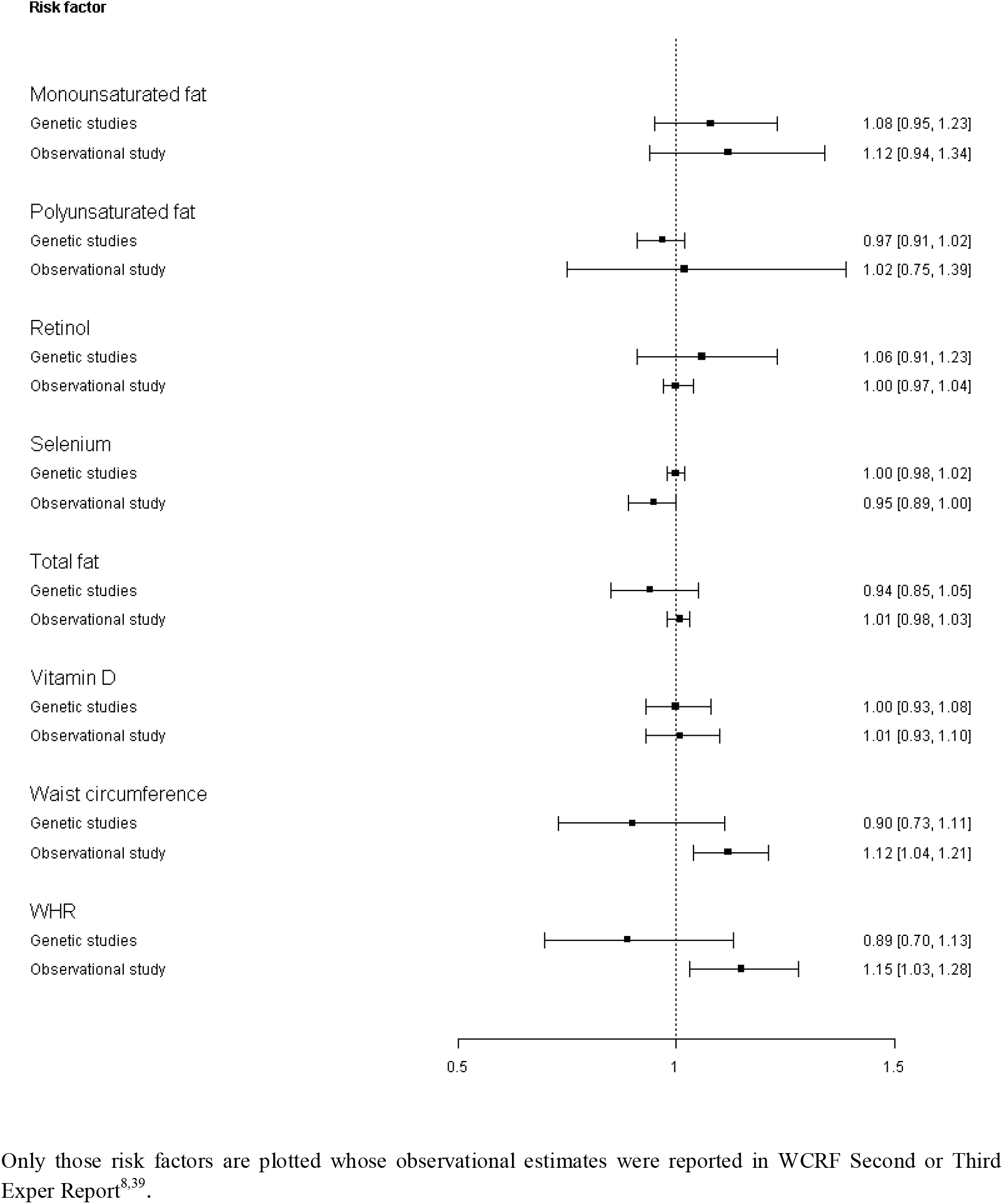
Comparison between observational and MR estimates for the risk factors and aggressive prostate cancer risk

There were only two SNPs available for the MR analysis of physical-activity (average acceleration) so we could not perform extensive sensitivity analyses. The direction of association for both SNPs was consistent (Supplementary File 1; Table S5) and the p-value for heterogeneity test was 0.99. These SNPs were on different chromosomes so represent independent associations. After exploring MRBASE-PheWAS (http://phewas.mrbase.org/), we found that these two SNPs were associated with anthropometric traits other than physical activity (Supplementary File 1; Table S6). The results for the effect of serum iron, and increasing BMI on overall prostate cancer were consistent across the various sensitivity analyses (Supplementary File 1; Figure S1-S3). The test for directional horizontal pleiotropy by MR-Egger (serum iron: intercept: 0.0005, *P=*0.97 and BMI: intercept: −0.0002, *P=*0.89) didn’t find evidence of pleiotropy. Mono-unsaturated fat showed results in the opposite direction using MR-Egger regression (OR: 0.85; 95% CI: 0.55, 1.31; *P*=0.51) compared to other MR-methods (Supplementary File 1; Figure S1 and S4). However, there was no strong evidence for directional pleiotropy for mono-unsaturated fat using MR-Egger (test for directional horizontal pleiotropy by MR-Egger: intercept: 0.04, *P=*0.31). The MR-Egger tests for both iron and mono-unsaturated fat had low power due to the small number of SNPs used, however all 5 SNPs for iron and 4/5 SNPs for mono-unsaturated fats showed associations with prostate cancer in the same direction (Supplementary File 1; Table S7-S8). At MRBASE-PheWAS, five SNPs of iron were associated with haemoglobin concentration and blood cells count (Supplementary File 1; Table S9) and SNPs of mono-unsaturated fat were associated with lipids (Supplementary File 1; Table S10). The MR results for single SNP analyses of BMI (overall prostate cancer) and height (aggressive prostate cancer) are provided in Supplementary File 1; Table S11-S12.

### Replication

The MR analyses were repeated using prostate cancer summary data generated from UK Biobank for physical-activity, iron, BMI, and mono-unsaturated fat (Figure 3). The point-estimates showed consistent directions of association for physical-activity (OR: 0.37; 95% CI: 0.13, 1.06; P=0.07), and BMI (OR: 0.83, 95% CI: 0.74, 0.94; P=0.002). The point-estimates for iron (OR: 1.07; 95% CI: 0.96, 1.20; P=0.24) and mono-unsaturated fat (OR: 0.89; 95% CI: 0.73, 1.07; P=0.20) were in the opposite direction, but the power to detect an effect with these risk factors in UK Biobank was low and confidence intervals for the replication analysis overlapped with our main analysis for all risk factors.

**Figure 3.**
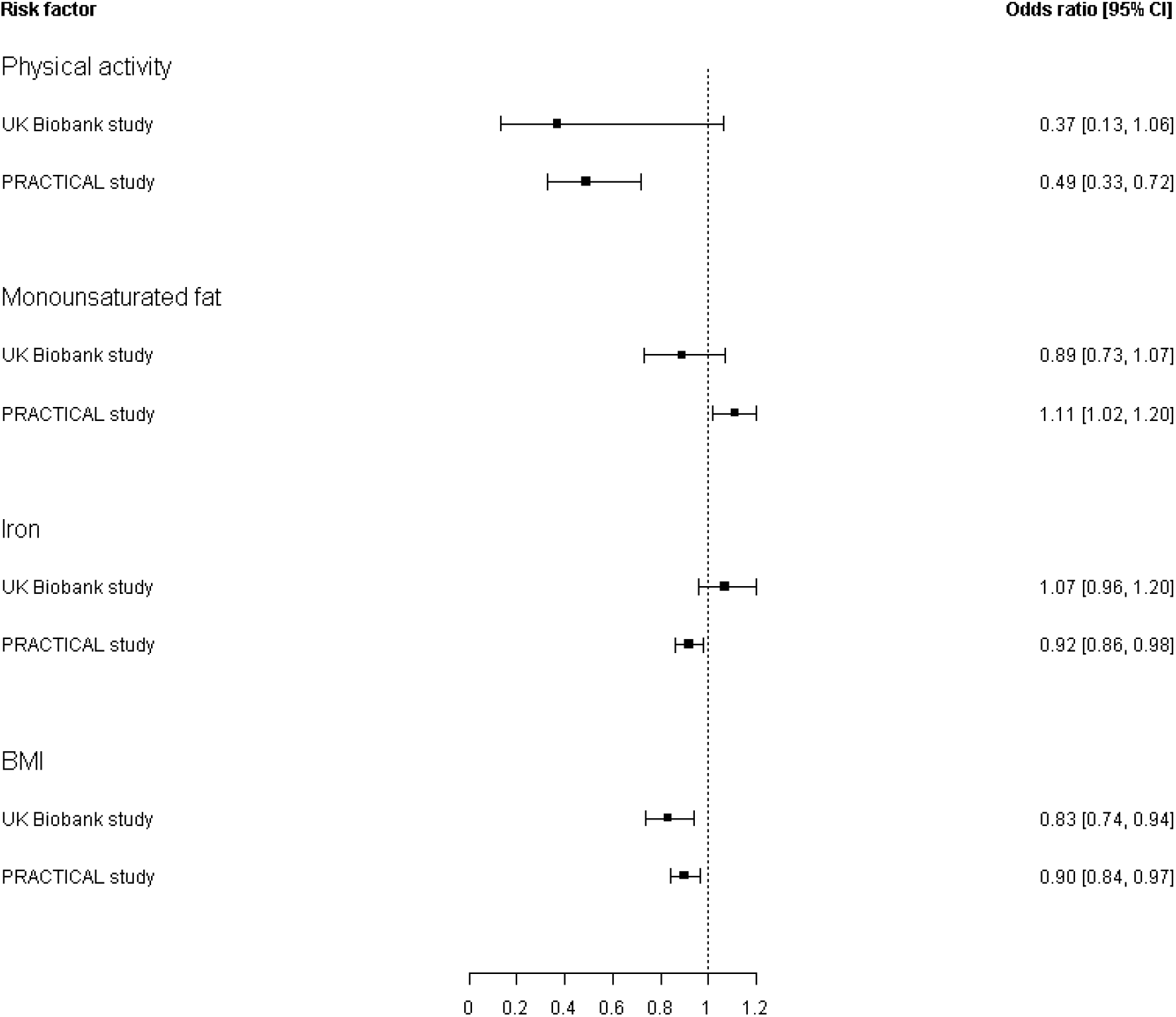
Comparison of MR estimates (OR) from the main and replication analyses for the risk factors that showed evidence of association (p<0.05) with overall prostate cancer in PRACTICAL

## Discussion

We found consistent evidence that physical-activity (assessed as ‘average accelerations’, but not other measures of physical activity), and BMI have an inverse effect on overall prostate cancer risk. There was also evidence of an inverse effect of iron and a positive effect of circulating mono-unsaturated fat in our initial analyses, the effect sizes were in opposite direction in UK Biobank study however power for the replication analysis was low and their CIs overlapped with the PRACTICAL study. There was weak evidence for physical-activity (average accelerations), and iron had a similar effect on aggressive prostate cancer to that seen for overall prostate cancer. We found little evidence that any of the other risk factors studied have a causal role in overall or aggressive prostate cancer, but height showed a positive association with aggressive prostate cancer. The CIs of MR results overlapped with those seen in the observational analyses for all risk factors except for average acceleration and mono-unsaturated fat (overall prostate cancer). In fact, confidence intervals for our MR analysis of aggressive prostate cancer were wide and the power for these analyses was low for many risk factors.

The WCRF report meta-analysed self-reported physical-activity which was assessed in different studies by various methods (i.e. occupational, recreational and total physical-activity) as highest versus lowest level of total physical-activity, a relatively crude dichotomy that may have masked associations. Our MR analysis which proxied fraction accelerations was the most similar to the observational analyses and we did not find evidence of an association of this measure with prostate cancer. However, we did find an association with average acceleration, which is a different measure and could be high if someone is consistently engaging in light-intensity activity across most of the waking day (vs lots of sitting and a 30 minutes bout of MVPA). Indeed, there is little genetic or phenotypic correlation between the two measures in the UKBiobank population^29^. The mechanism for our association between average accelerations and prostate cancer is unclear, although this could be through improved insulin sensitivity or reduced insulin-like growth factor-1 (IGF-1)^30^, reduced levels of testosterone and dihydrotestosterone^31^, alterations in the antioxidant defence system^31^, or improvements in the immune system through enhanced natural killer cell activity^31^.

Whilst not consistent with observational analyses, our results for BMI are concordant with other lines of evidence. We have previously shown weak evidence that higher BMI is associated with a reduced prostate cancer risk, in a smaller sample from the PRACTICAL consortium^32^. A study examined childhood and adult body size in relation to total incident prostate cancer in a prospective cohort of 47,491 US men^33^. High BMI at age 21 was inversely associated with total prostate cancer, with fatal and advanced disease. The association for late adult BMI was more complex and differed by age this could represent confounding by other factors for example physical activity, and diet etc.

Despite showing a protective effect of BMI on prostate cancer risk, we did not find any strong evidence of waist circumference or WHR with risk. However, the results for all these risk factors were in the same direction. The point estimates for BMI, waist circumference and WHR were in the opposite direction to findings from observational results for aggressive prostate cancer. For aggressive prostate cancer, power calculations suggested that we would have good power to detect an effect of BMI (96%) but very low power (33% and 23%) to detect an OR of 1.20 (or, conversely a protective OR of at least 0.80) for waist circumference and WHR respectively. An increased estrogen production has been observed in obese men^34^. The sensitivity of prostate cancer to sex hormones has been exploited for therapeutic purposes for many years. Androgen-deprivation therapy is a common treatment in prostate cancer^35^, as is the therapeutic use of estrogen for patients with metastatic prostate cancer in both the US^36^ and Europe^37^, though not popular due to cardiovascular and other side effects. Hence the clinical prediction would be that obesity would be associated with a lower risk of prostate cancer. However, we cannot rule out detection bias^38^ arising from delayed diagnosis and therefore a more advanced stage at diagnosis in obese men. This could arise due to lower accuracy of digital rectal examination in obese men or lower PSA values caused by obesity-related traits.

The analysis of iron as a risk factor for prostate cancer found that the evidence was too limited to draw conclusions in the WCRF Second Expert Report and this was not updated in the Third Expert Report due to a lack of new evidence^8,39^. Population studies that have examined the associations between serum iron and cancer outcomes are limited and have reported discordant findings. A prospective cohort study with 15-16 years follow up time reported higher serum iron concentrations increased non-skin cancer risk overall but conversely, in men, higher serum iron concentrations decreased the risk of non-skin cancer^40^. A Swedish cohort reported increased serum iron concentrations were not associated with overall cancer risk except for a slightly higher risk of postmenopausal breast cancer^41^. Previous findings have suggested that, although higher circulating iron concentrations may potentially increase the risk of cancer in women it may be protective against cancer in men^40^ which is in accordance with our MR findings. Although again we were unable to replicate these findings in UK Biobank due to low power.

Our MR-analysis investigated the association of circulating levels of mono-unsaturated fat on prostate cancer, circulating levels of this nutrient have been shown to be poorly correlated with mono-unsaturated fat intake measured by questionnaire^42^. The inclusion of an objective measure in our MR-analysis versus questionnaire data in observational studies could be the reason for the discordant results. Although discordance could also be due to negative confounding in the former studies resulting in the effect estimate being closer to the null (e.g., through other dietary, lifestyle, or molecular factors. Instruments for mono-unsaturated fat are correlated with instruments for saturated fat and other lipids. It could be possible that saturated fat or other lipids cause prostate cancer and this result might reflect that. More research is needed into the independence of mono-unsaturated fat of other lipid factors.

Our MR results showed positive association between height and aggressive prostate cancer and the results was consistent with observational studies. Adult height is associated with the rate of growth during fetal life and childhood^43,44^.Health and nutrition status in the neonatal period and childhood may affect the age of sexual maturity. These processes are mediated by changes in the hormonal microenvironment that may have both short and long term effects on circulating levels of growth factors, insulin and other endocrine or tissue specific mediators that may impact cancer risk^45^.

The results from replication analyses were compatible with the main findings for BMI, and physical activity (albeit with wide confidence intervals for physical activity), which increases the likelihood that these findings are real. The UK Biobank sample size was however, smaller, and the effects were estimated with less precision and whilst the results for iron and mono-unsaturated fats did not appear to replicate, due to low power in the replication study we cannot rule out causal effects of these nutrients.

This study’s major strength is the use of MR, which is less susceptible to problems of measurement error, confounding and reverse causation in comparison to conventional observational studies. The use of two-sample MR enabled the use of the largest GWAS of prostate cancer^21^ to date. We were also able to make use of the largest GWASs on the risk factors of interest, to increase the precision of the SNP-exposure estimates, which should reduce impact of weak instruments bias, which in turn increase statistical power assuming the SNP-exposure estimates are unbiased and risk factor/outcome samples come from the same population.

The study also has some limitations. We had only two SNPs for physical-activity assessed as average acceleration. If there were many independent SNPs available the causal inference could have been strengthened because a) each variant represents an independent natural experiment, and a more precise overall causal estimate (i.e. tighter CIs) can be obtained by meta-analysing the single estimates from each instrument; and b)potential bias arising from the violation of the assumptions can be detected or corrected by evaluating the consistency of effects across instruments^16,24,28,46,47^. For many of the risk factors reported in the WCRF 2018 report, for example alpha-tocopheral, vitamin A, vitamin C etc, we did not find genetic instruments to conduct MR analyses. For the majority of the risk factors in overall prostate cancer, MR analyses were sufficiently powered to detect effect sizes of a modest magnitude (OR of 1.20 or 0.80) except for physical-activity traits (overall acceleration average, fraction of accelerations >425 milli-gravities, and sedentary behaviour), thus failure to detect strong evidence of effects for these risk factors could be due to low power to detect smaller effect sizes. Further identification of independent genetic variants that influence these risk factors will help to improve statistical power for future analyses.

In conclusion, we found evidence that physical-activity, serum iron, and BMI may be causally and inversely related to and circulating mono-unsaturated fat and height may be causally and positively related to, prostate cancer risk. Further studies should investigate the mechanisms by which these factors may lead to prostate cancer and investigate the potential to intervene to reduce risk.

## Contributors

SL, RMM and NK conceived and designed the study. NK conducted the analyses. NK wrote the manuscript with input from all authors. Correspondence and material requests should be addressed to SL.

## Acknowledgments

This research was funded by a grant awarded to SL for 3 years to identify modifiable risk factors for prostate cancer, by the World Cancer Research Fund International (Grant Reference Number: 2015/1421), a Cancer Research UK program grant (C18281/A19169) and the National Institute for Health Research (NIHR) Bristol Biomedical Research Centre. Dr Haycock is supported by CRUK Population Research Postdoctoral Fellowship C52724/A20138. The views expressed are those of the author(s) and not necessarily those of the NIHR or the Department of Health and Social Care. Lynch was supported by a Mid-Career Research Fellowship from the Victorian Cancer Agency (MCRF18005).

