## Supplemental material for "Appraising causal relationships of dietary, nutritional and physical-activity exposures with overall and aggressive prostate cancer: two-sample Mendelian randomization study based on 79,148 prostate cancer cases and 61,106 controls"

### Defining genetic instruments

For the risk factors with GWAS data, we noted the PubMed ID of the GWAS used, sample size, total number of GWAS significant SNPs and their rs numbers, units of the analysis and the population standard deviation (SD) for each risk factor. We also extracted SNP-trait summary data directly from the published articles. When there were multiple GWAS results for the same risk factor, the largest GWAS of that risk factor was used where possible to maximise the precision of the instrument and the statistical power of the analyses. If the largest GWAS didn’t yield any SNPs which we could use in our analysis (e.g. due to missing data in the outcome dataset or difficulty harmonising^1^ to the same reference allele for both exposure and outcome) then the second largest GWAS was used, and so on.

After selecting the set of SNPs for each risk factor *(*which passed a conventional GWAS-significant *P*-value threshold of <5x10^-08^), we extracted the following information for each SNP-risk factor association: effect allele, other allele, beta coefficient and standard error (SE). We removed any SNP that was missing this information. We converted each beta-coefficient and corresponding SE reported in the original GWAS to SD units by dividing them by the SD of the sample mean of the trait in the study population where necessary and possible (e.g. some were already in SD units). When the beta-coefficient and corresponding SE were reported in logarithmic scale in the original GWAS and the Mean and SD were reported on the original scale, we converted the SD unit to the logarithm scale using the following equations:

$$variance=log10(\left( {SD}^{2}+ {Mean}^{2} \right)/{SD}^{2})$$

$$SD= \surd variance$$

We then divided the beta-coefficients and corresponding SE by the SD which is also on logarithmic scale. Risk factors were ‘instrumentable’ if a relevant GWAS was available and SNPs required to generate reliable instruments did not have missing information.

For the SNP(s) extracted for use in the MR-analysis, we calculated the proportion of variance explained (R^2^) in the risk factor by the SNP(s) and the strength of the instrument (F-statistic)^2^. The formulas to calculate R^2^ and F-statistic were:

$R^{2}= (2\times\beta^{2}\times MAF\times(1-MAF))/(2 \times\beta^{2}\times MAF\times(1-MAF)+{(SE\left( \beta\right))}^{2} \times(2 \times N)\times MAF\times(1-MAF))$, where β is the effect size (beta coefficient) for a given SNP, MAF is the minor allele frequency, SE(β) is the standard error of the effect size, and N is the sample size of the GWAS for the SNP-risk factor association.

$F=R^{2}\times(N-1-k))/((1-R^{2} ) \times k)$, where R^2^ is the proportion of variance explained in the risk factor by the genetic instrument, N is the sample size of the GWAS, k is the number of SNPs included in the instrument.

Once instruments have been identified for the risk factor, the European samples from the 1000 genomes project were used to estimate linkage disequilibrium (LD) between SNPs. The SNPs were extracted from 1000 genomes data, correlations calculated between them, and amongst those SNPs that had r^2^ >0.01 (assessed within 10,000 kilobase pair windows) only the SNP with the lowest P-value for the association with the risk factor of interest was retained. Exposure and outcome data were harmonized such that the effect of each SNP on the outcome and exposure was relative to the same allele. For palindromic SNPs (i.e., those where the effect/other allele were either an A/T or G/C combination), we used allele frequency to resolve strand ambiguity and discarded palindromic SNP(s) that had minor allele frequency above 0.42^1^.

We calculated the power to detect an odds ratio (OR) of 1.2 (or conversely a protective OR of 0.80) for each risk factor given an alpha-level of 0.05, the variance explained in the risk factor by the instrument and the sample size (79,148 prostate cancer cases and 61,106 controls), as described peviously^3^.

There were several GWASs of physical activity available that were based on the UK Biobank population^4,5^. We included only those that estimated physical activity or movement related behaviours using accelerometer data^6^ (subject to less measurement error than self-reported activity), and used four summary measures: overall acceleration average, fraction of accelerations >425 milli-gravities (mg) (with total activity as the denominator), sedentary behaviour and sleep duration, as previously described^4,6^. A previous study derived physical activity data from Axivity AX3 triaxial accelerometers, worn on the wrist for a seven-day protocol^6^. 100Hz raw triaxial acceleration data was used after calibration, removal of gravity and sensor noise, and identification of wear/non-wear episodes. Non-wear time was defined as consecutive stationary episodes lasting for at least 60 minutes where all three axes had a standard deviation of less than 13.0 mg. For overall activity levels, average vector magnitude for each 30 seconds epoch was selected, which is the recommended variable for activity analysis^24^. To predict sleep and sedentary behaviour, the study used a machine-learning method (balanced random forests with Markov confusion matrices) to predict behaviour for each 30 second epoch in the participants’ accelerometer data^4^. Sedentary behaviour was defined as one which had a metabolic equivalent of task (MET) energy expenditure score of ≤ 1.5 that occurred in a sitting, lying, or reclining posture^4^.

For nutritional factors, we only included GWASs of circulating levels of macro- and micro-nutrients measured in blood. In a meta-analysis of iron reported previously, most studies measured serum iron using colorimetric assay^7^. Serum phosphorus concentrations were quantified using an automated platform in which inorganic phosphorus reacts with ammonium molybdate in an acidic solution to form a colored phosphomolybdate complex^8^. Reverse-phase high-pressure liquid chromatography was utilized to assess serum alpha-carotene, beta-carotene, lycopene and retinol concentrations^9-12^. Most studies that participated in a meta-analysis of calcium measured it using a colorimetric assay^13^. The SNPs to proxy for selenium was constructed by meta-analysing blood and toenail selenium GWAS^14^. 25-hydroxyvitamin D was measured with high-performance liquid chromatography-tandem mass spectrometry^15^.

Fasting glucose was previously measured in mM. The study excluded individuals from the analysis if they had a physician diagnosis of diabetes, were on diabetes treatment (oral or insulin) or had a fasting plasma glucose concentration equal to or greater than 7 mM^16^. A previous study used high-throughput NMR metabolomics platform to quantify human blood metabolites including mono-unsaturated fat, polyunsaturated fat and total fat^17^. All dietary factors were self-reported (see main Table 1).

Birth weight was collected from a variety of sources, including measurements at birth by medical practitioners, obstetric records, medical registers, interviews with the mother and self-report as adults^18^. Body mass index (BMI) value was constructed from height and weight. BMI, height, waist circumference and waist-hip ratio was either measured at clinic or self-reported^19-21^.

**Sensitivity analyses**

There was evidence of heterogeneity for several risk factors (P_het_ < 0.05; calculated by Cochran’s Q test) for both overall and aggressive prostate cancer risk (Table S3 and Table S4). However, the results of sensitivity analyses for these risk factors were largely consistent with the main IVW analyses (Figure S1 and S6). MR-PRESSO revealed outlying SNPs for birth weight, BMI, height, waist circumference, WHR, glucose, total fat, and alcohol. After removing the outlying SNPs, the IVW analysis produced similar results for all these risk factors compared to results with all SNPs (overall prostate cancer risk, Figure S1). The outlying SNPs for each risk factor are provided in Table S13 and Table S14 for overall prostate cancer and aggressive prostate cancer respectively. There was weak evidence of protective association between waist circumference and overall prostate cancer risk after removing the outlying SNPs (OR: 0.89; 95% CI: 0.79, 1.00; P=0.06) and the direction of association was consistent across all other analyses. For aggressive prostate cancer risk, MR-PRESSO revealed outlying SNPs for birth weight, BMI, height, waist circumference and alcohol. After removing the outlying SNPs, the IVW analysis produced similar results for all these risk factors (Figure S6).

**Table S1. Observational associations of previously reported risk factors for overall prostate cancer risk^22^*.***

| **Exposure** | **N cases** | **RR (95% CI)** | **Units** | ***P*-value** |
| --- | --- | --- | --- | --- |
| ***Anthropometrics and other measures*** | | | | |
| Birth weight | 2827 | 1.03 (0.99, 1.08) | 500 g | 0.18 |
| BMI | 91486 | 1.00 (0.98, 1.03) | 5 kg/m^2^ | 1.00 |
| Height | 79387 | 1.04 (1.03, 1.05) | 5 cm | 1.3x10^-15^ |
| Waist circumference | 6883 | 1.00 (0.97, 1.03) | 10 cm | 1.00 |
| WHR | 5843 | 1.01 (0.96, 1.06) | 0.1 units | 0.69 |
| **C*irculating macro- and micro-nutrients*** | | | | |
| Sugar/Sucrose* | N/A | N/A | N/A | N/A |
| Monounsaturated fat | 4384 | 1.00 (0.99, 1.01) | 10 g/day | 1.00 |
| Polyunsaturated fat | 4766 | 1.00 (0.93, 1.08) | 10 g/day | 1.00 |
| Total fat | 6063 | 1.00 (0.99, 1.01) | 10 g/day | 1.00 |
| Alpha-carotene | 2833 | 1.06 (0.94, 1.21) | 10 mcg/100 ml | 0.37 |
| Beta-carotene | 3449 | 0.99 (0.95, 1.04) | 10 mcg/100 ml | 0.66 |
| Calcium | 38749 | 1.05 (1.02, 1.09) | 400 mg per day | 0.004 |
| Iron* | N/A | N/A | N/A | N/A |
| Lycopene | 4665 | 0.99 (0.96, 1.01) | 10 mcg/dl | 0.44 |
| Phosphsorous* | N/A | N/A | N/A | N/A |
| Retinol | 7168 | 1.01 (1.00, 1.03) | 10 mcg/100 ml | 0.19 |
| Selenium | 3559 | 0.95 (0.91, 1.00) | 10 mcg/l | 0.03 |
| Vitamin D | 7802 | 1.04 (1.00, 1.07) | 30 nmol/l | 0.02 |
| ***Consumption of foods and drinks*** | | | | |
| Alcohol | 36942 | 1.01 (0.99, 1.02) | 1 drink/day | 0.19 |
| Coffee | 9841 | 0.99 (0.98, 1.00) | 1 cup/day | 0.05 |
| Dairy products | 38107 | 1.07 (1.02, 1.12) | 400 g/d | 0.005 |
| ***Physical activity*** | | | | |
| Physical activity | 23478 | 0.97 (0.9, 1.04) | Highest versus lowest total physical activity | 0.41 |

BMI = body mass index; RR = risk ratio; WHR = waist-hip ratio

* The WCRF found limited evidence of limited-no-conclusion for these risk factors and did not publish the observational results in Second or Third Exper Report^22,23^.

N/A = not accessible/available

**Table S2. Observational associations of previously reported risk factors for aggressive prostate cancer risk^22^*.***

| **Exposure** | **Studies** | **RR (95% CI)** | **Units** | ***P*-value** |
| --- | --- | --- | --- | --- |
| ***Anthropometrics and other measures*** | | | | |
| Birth weight | 2 | 1.09 (0.97, 1.22) | 500 g | 0.14 |
| BMI | 23 | 1.08 (1.04, 1.12) | 5 kg/m^2^ | 4.68×10^-5^ |
| Height | 19 | 1.04 (1.02, 1.06) | 5 cm | 6.4×10^-5^ |
| Waist circumference | 4 | 1.12 (1.04, 1.21) | 10 cm | 0.003 |
| WHR | 4 | 1.15 (1.03, 1.28) | 0.1 units | 0.01 |
| **C*irculating macro- and micro-nutrients*** | | | | |
| Sugar/Sucrose* | N/A | N/A | N/A | N/A |
| Monounsaturated fat | 4 | 1.12 (0.94, 1.34) | 10 g/day | 0.21 |
| Polyunsaturated fat | 4 | 1.02 (0.75, 1.39) | 10 g/day | 0.90 |
| Total fat | 3 | 1.01 (0.98, 1.03) | 10 g/day | 0.43 |
| Alpha-carotene* | N/A | N/A | N/A | N/A |
| Beta-carotene | 3 | 0.97 (0.85, 1.12) | 10 mcg/100 ml | 0.67 |
| Calcium | 10 | 1.02 (0.93, 1.12) | 400 mg per day | 0.67 |
| Iron* | N/A | N/A | N/A | N/A |
| Lycopene | 5 | 0.98 (0.93, 1.03) | 10 mcg/dl | 0.44 |
| Phosphorous* | N/A | N/A | N/A | N/A |
| Retinol | 4 | 1.00 (0.97, 1.04) | 10 mcg/100 ml | 1.00 |
| Selenium | 5 | 0.95 (0.89, 1.00) | 10 mcg/l | 0.08 |
| Vitamin D | 6 | 1.01 (0.93, 1.10) | 30 nmol/l | 0.82 |
| ***Consumption of foods and drinks*** | | | | |
| Alcohol | 7 | 1.00 (0.96, 1.03) | 1 drink/day | 1.00 |
| Coffee | N/A | N/A | N/A | N/A |
| Dairy products | 10 | 0.97 (0.91, 1.05) | 400 g/d | 0.40 |
| ***Physical activity*** | | | | |
| Physical activity | N/A | N/A | N/A | N/A |

BMI = body mass index; RR = risk ratio; WHR = waist-hip ratio

* The WCRF found limited evidence of limited-no-conclusion for these risk factors and did not publish the observational results in Second or Third Exper Report^22,23^.

N/A = not accessible/available

**Table S3. Mendelian randomization analyses of the association between risk factors and overall prostate cancer risk**

| **Risk factor** | **# SNPs** | **MR method** | **OR** | **LCI** | **UCI** | **P_assoc_** | **P_pltr_** | **P_het_** |
| --- | --- | --- | --- | --- | --- | --- | --- | --- |
| ***Anthropometrics and other measures*** | | | | | | | | |
| Birth weight | 46 | IVW random | 1.03 | 0.89 | 1.18 | 0.73 | 0.84 | 3.71×10^-8^ |
| BMI | 535 | IVW random | 0.90 | 0.84 | 0.97 | 0.003 | 0.89 | 1.19×10^-41^ |
| Height | 433 | IVW random | 1.02 | 0.97 | 1.07 | 0.50 | 0.10 | 2.33×10^-51^ |
| Waist circumference | 45 | IVW random | 0.91 | 0.77 | 1.09 | 0.32 | 0.88 | 8.78×10^-12^ |
| WHR | 31 | IVW random | 0.92 | 0.78 | 1.08 | 0.30 | 0.50 | 0.007 |
| **C*irculating macro- and micro-nutrients*** | | | | | | | | |
| Sugar/Sucrose (fasting glucose) | 15 | IVW random | 0.89 | 0.62 | 1.28 | 0.54 | 0.22 | 9.08×10^-5^ |
| Monounsaturated fat | 5 | IVW random | 1.11 | 1.02 | 1.20 | 0.02 | 0.31 | 0.31 |
| Polyunsaturated fat | 19 | IVW random | 1.01 | 0.97 | 1.06 | 0.50 | 0.47 | 0.03 |
| Total fat | 12 | IVW random | 0.99 | 0.91 | 1.08 | 0.84 | 0.30 | 0.005 |
| Alpha-carotene | 3 | IVW random | 1.00 | 0.98 | 1.03 | 0.72 | 0.67 | 0.07 |
| Beta-carotene | 1 | Wald ratio | 0.97 | 0.90 | 1.04 | 0.37 | - | - |
| Calcium | 5 | IVW fixed | 0.66 | 0.38 | 1.14 | 0.14 | 0.51 | 0.73 |
| Iron | 5 | IVW random | 0.92 | 0.86 | 0.98 | 0.007 | 0.97 | 0.30 |
| Lycopene | 1 | Wald ratio | 0.97 | 0.88 | 1.07 | 0.55 | - | - |
| Phosphorous | 4 | IVW random | 1.03 | 0.90 | 1.18 | 0.69 | 0.25 | 0.18 |
| Retinol | 2 | IVW fixed | 1.01 | 0.93 | 1.10 | 0.83 | - | 0.41 |
| Selenium | 2 | IVW fixed | 1.00 | 0.98 | 1.01 | 0.47 | - | 0.61 |
| Vitamin D | 4 | IVW fixed | 1.00 | 0.97 | 1.03 | 0.90 | 0.47 | 0.52 |
| ***Consumption of foods and drinks*** | | | | | | | | |
| Alcohol | 77 | IVW random | 0.96 | 0.72 | 1.28 | 0.77 | 0.43 | 1.41×10^-22^ |
| Coffee | 4 | IVW random | 1.16 | 0.86 | 1.56 | 0.32 | 0.94 | 0.02 |
| Dairy products (milk intake) | 1 | Wald ratio | 1.00 | 0.97 | 1.03 | 0.92 | - | - |
| ***Physical activity*** | | | | | | | | |
| Average acceleration | 2 | IVW fixed | 0.49 | 0.33 | 0.72 | 0.0003 | - | 0.99 |
| Sedentary behaviour | 2 | IVW fixed | 0.95 | 0.63 | 1.42 | 0.79 | - | 0.54 |
| Fraction accelerations | 1 | Wald ratio | 1.06 | 0.56 | 2.02 | 0.85 | - | - |
| Sleep duration | 7 | IVW random | 1.00 | 0.80 | 1.24 | 0.97 | 0.85 | 0.20 |

BMI = body mass index; LCI = lower 95% confidence interval; OR = odds ratio; P_assoc_ = P-value for association; P_het_ = P-value for heterogeneity between instrumental SNP causal estimates; P_pltr_ = P-value for horizontal pleiotropy from MR-Egger intercept test; WHR = waist-hip ratio; UCI = upper 95% confidence interval

\# SNPs represents the number of SNPs used within the instrument for each risk factor after clumping, harmonization and extraction from data of overall prostate cancer

**Table S4. Mendelian randomization analyses of the association between risk factors and aggressive prostate cancer risk**

| **Risk factor** | **# SNPs** | **MR method** | **OR** | **LCI** | **UCI** | **P_assoc_** | **P_pltr_** | **P_het_** |
| --- | --- | --- | --- | --- | --- | --- | --- | --- |
| ***Anthropometrics and other measures*** | | | | | | | | |
| Birth weight | 45 | IVW random | 1.12 | 0.92 | 1.35 | 0.26 | 0.42 | 0.01 |
| BMI | 525 | IVW random | 0.94 | 0.85 | 1.03 | 0.19 | 0.76 | 1.97×10^-8^ |
| Height | 426 | IVW random | 1.07 | 1.01 | 1.15 | 0.03 | 0.54 | 1.28×10^-7^ |
| Waist circumference | 44 | IVW random | 0.90 | 0.73 | 1.11 | 0.34 | 0.63 | 0.02 |
| WHR | 28 | IVW random | 0.89 | 0.70 | 1.13 | 0.33 | 0.66 | 0.37 |
| **C*irculating macro- and micro-nutrients*** | | | | | | | | |
| Sugar/Sucrose (glucose) | 15 | IVW random | 0.90 | 0.62 | 1.30 | 0.56 | 0.63 | 0.38 |
| Monounsaturated fat | 5 | IVW fixed | 1.08 | 0.95 | 1.23 | 0.26 | 0.43 | 0.42 |
| Polyunsaturated fat | 19 | IVW random | 0.97 | 0.91 | 1.02 | 0.24 | 0.38 | 0.28 |
| Total fat | 12 | IVW random | 0.94 | 0.85 | 1.05 | 0.28 | 0.75 | 0.20 |
| Alpha-carotene | 3 | IVW fixed | 1.01 | 0.98 | 1.03 | 0.55 | 0.99 | 0.73 |
| Beta-carotene | 1 | Wald ratio | 0.94 | 0.83 | 1.06 | 0.31 | N/A | N/A |
| Calcium | 5 | IVW fixed | 0.94 | 0.36 | 2.42 | 0.89 | 0.93 | 0.90 |
| Iron | 5 | IVW fixed | 0.92 | 0.84 | 1.02 | 0.10 | 0.76 | 0.66 |
| Lycopene | 1 | Wald ratio | 0.96 | 0.81 | 1.14 | 0.62 | N/A | N/A |
| Phosphorous | 4 | IVW random | 1.09 | 0.85 | 1.41 | 0.50 | 0.60 | 0.11 |
| Retinol | 2 | IVW fixed | 1.06 | 0.91 | 1.23 | 0.44 | N/A | 0.70 |
| Selenium | 2 | IVW fixed | 1.00 | 0.98 | 1.02 | 0.97 | N/A | 0.42 |
| Vitamin D | 4 | IVW random | 1.00 | 0.93 | 1.08 | 0.95 | 0.47 | 0.15 |
| ***Consumption of foods and drinks*** | | | | | | | | |
| Alcohol | 76 | IVW random | 0.94 | 0.66 | 1.35 | 0.75 | 0.47 | 4.49×10^-5^ |
| Coffee | 4 | IVW random | 1.22 | 0.88 | 1.69 | 0.24 | 0.88 | 0.24 |
| Dairy products (milk intake) | 1 | Wald ratio | 0.99 | 0.94 | 1.05 | 0.81 | N/A | N/A |
| ***Physical activity*** | | | | | | | | |
| Average acceleration | 2 | IVW fixed | 0.51 | 0.26 | 1.01 | 0.05 | N/A | 0.88 |
| Sedentary behaviour | 2 | IVW fixed | 0.59 | 0.15 | 2.41 | 0.46 | N/A | 0.04 |
| Fraction accelerations | 1 | Wald ratio | 1.67 | 0.56 | 5.01 | 0.36 | N/A | N/A |
| Sleep duration | 7 | IVW fixed | 1.00 | 0.73 | 1.38 | 0.98 | 0.25 | 0.48 |

BMI = body mass index; LCI = lower 95% confidence interval; OR = odds ratio; P_assoc_ = P-value for association; P_het_ = P-value for heterogeneity between instrumental SNP causal estimates; P_pltr_ = P-value for horizontal pleiotropy from MR-Egger intercept test; WHR = waist-hip ratio; UCI = upper 95% confidence interval

\# SNPs represents the number of SNPs used within the instrument for each risk factor after clumping, harmonization and extraction from data of overall prostate cancer

**Table S5: The MR association of individual SNPs of average acceleration with overall prostate cancer risk**

| **SNP** | **OR** | **LCI** | **UCI** | **P_assoc_** |
| --- | --- | --- | --- | --- |
| rs55657917 | 0.49 | 0.29 | 0.82 | 0.007 |
| rs59499656 | 0.49 | 0.27 | 0.88 | 0.02 |

**Table S6. Evidence of association (p<5×10^-8^) of average acceleration SNPs with other traits**

| **SNP** | **CHR** | **Gene** | **Trait** |
| --- | --- | --- | --- |
| rs55657917 | 17 | CRHR1 | Alcohol intake frequency (UK biobank)  Height (UK biobank)  Ovarian cancer (PMID: 28346442)  Red blood cell count (PMID: 27863252)  Systolic blood pressure (UK biobank)  Puberty (UK biobank)  Anxiety (UK biobank) |
| rs59499656 | 18 | RIT2/SYT4 | Adiposity (UK biobank)  ΒΜΙ (UK biobank)  Weight (UK biobank) |

**Table S7. The MR association of individual SNPs of iron with overall prostate cancer risk**

| **SNP** | **OR** | **LCI** | **UCI** | **P_assoc_** |
| --- | --- | --- | --- | --- |
| rs1799945 | 0.84 | 0.75 | 0.95 | 0.004 |
| rs1800562 | 0.91 | 0.82 | 1.01 | 0.07 |
| rs7385804 | 0.84 | 0.64 | 1.08 | 0.18 |
| rs8177240 | 0.99 | 0.77 | 1.28 | 0.96 |
| rs855791 | 0.98 | 0.89 | 1.07 | 0.61 |

**Table S8. The MR association of individual SNPs of mono-unsaturated fat with overall prostate cancer risk**

| **SNP** | **OR** | **LCI** | **UCI** | **P_assoc_** |
| --- | --- | --- | --- | --- |
| rs115849089 | 1.06 | 0.86 | 1.30 | 0.60 |
| rs1260326 | 1.21 | 1.05 | 1.38 | 0.01 |
| rs1800588 | 1.15 | 0.99 | 1.33 | 0.07 |
| rs41272659 | 1.05 | 0.84 | 1.30 | 0.69 |
| rs8107974 | 0.94 | 0.78 | 1.15 | 0.57 |

**Table S9. Evidence of association (p<5×10^-8^) of iron SNPs with other traits**

| **SNP** | **CHR** | **Gene** | **Trait** |
| --- | --- | --- | --- |
| rs1799945 | 6 | HFE | Haemoglobin (PMID: 27863252)  Blood cell count (PMID: 27863252)  Blood pressure (UK biobank) |
| rs1800562 | 6 | HFE | Disorders of iron metabolism (UK biobank)  Haemoglobin (PMID: 27863252)  Ferritin (PMID: 25352340)  Blood cell count (PMID: 27863252)  LDL cholesterol, total cholesterol (PMID: 24097068) |
| rs7385804 | 7 | TFR2 | Haemoglobin (PMID: 27863252)  Blood cell count (PMID: 27863252) |
| rs8177240 | 3 | TF | Transferrin (PMID: 25352340)  Haemoglobin (PMID: 27863252) |
| rs855791 | 22 | TMPRSS6 | Haemoglobin (PMID: 27863252)  Blood cell count (PMID: 27863252) |

**Table S10. Evidence of association (p<5×10^-8^) of mono-unsaturated fat SNPs with other traits**

| **SNP** | **CHR** | **Gene** | **Trait** |
| --- | --- | --- | --- |
| rs115849089 | 8 | LPL | Triglycerides (PMID: 27005778)  Lipids (PMID: 27005778)  Total cholesterol (PMID: 27005778)  Lipoprotein (PMID: 27005778)  Glycoprotein acetyls (PMID: 27005778)  Omega-7, omega-9 and saturated fatty acids (PMID: 27005778) |
| rs1260326 | 2 | GCKR | Triglycerides (PMID: 24097068)  Alcohol (PMID: 28937693 and UK biobank)  Adiposity (UK biobank)  Total cholesterol (PMID: 24097068)  Granulocyte percentage of myeloid (PMID: 27863252)  Fasting glucose (PMID: 22581228)  Triglycerides (PMID: 27005778)  Lipoprotein (PMID: 27005778)  Blood cell count (PMID: 27863252)  Omega-6 (PMID: 27005778)  Type 2 diabetes (PMID: 26551672) |
| rs1800588 | 15 | LIPC/ LOC101928635 | Triglycerides (PMID: 27005778)  HDL cholesterol (PMID: 24097068)  Cholesterol (PMID: 27005778)  Omega- 6, Omega-7, omega-9 and saturated fatty acids (PMID: 27005778) |
| rs41272659 | 2 | FASTKD2 | None |
| rs8107974 | 19 | SUGP1 | Cholesterol (PMID: 27005778)  Triglycerides (PMID: 27005778)  Lipids (PMID: 27005778)  Adiposity (UK biobank)  Omega-3, Omega-7, omega-9 and saturated fatty acids |

**Table S11: The MR association of individual SNPs of BMI with overall prostate cancer risk**

| **SNP** | **OR** | **LCI** | **UCI** | **P_assoc_** |
| --- | --- | --- | --- | --- |
| rs2185027 | 0.01 | 0.002 | 0.02 | 4.14×10^-15^ |
| rs11629783 | 0.02 | 0.01 | 0.07 | 1.10×10^-9^ |
| rs905938 | 0.03 | 0.01 | 0.10 | 8.37×10^-9^ |
| rs3826705 | 0.06 | 0.01 | 0.30 | 5.54×10^-4^ |
| rs3829849 | 0.07 | 0.01 | 0.38 | 0.002 |
| rs769674 | 0.07 | 0.02 | 0.26 | 5.04×10^-5^ |
| rs892261 | 0.07 | 0.02 | 0.33 | 7.51×10^-4^ |
| rs10971712 | 0.08 | 0.02 | 0.29 | 7.52×10^-5^ |
| rs11781699 | 0.08 | 0.02 | 0.40 | 0.002 |
| rs4278019 | 0.10 | 0.02 | 0.46 | 0.003 |
| rs329124 | 0.11 | 0.03 | 0.36 | 2.89×10^-4^ |
| rs7730004 | 0.12 | 0.04 | 0.38 | 2.91×10^-4^ |
| rs799449 | 0.13 | 0.04 | 0.46 | 0.002 |
| rs9615905 | 0.13 | 0.03 | 0.59 | 0.01 |
| rs1787267 | 0.14 | 0.04 | 0.55 | 0.005 |
| rs1865341 | 0.14 | 0.03 | 0.59 | 0.01 |
| rs2075205 | 0.14 | 0.03 | 0.60 | 0.01 |
| rs2866816 | 0.14 | 0.03 | 0.60 | 0.01 |
| rs329651 | 0.14 | 0.04 | 0.49 | 0.002 |
| rs3850422 | 0.14 | 0.03 | 0.56 | 0.01 |
| rs5396 | 0.14 | 0.04 | 0.46 | 0.001 |
| rs2293605 | 0.15 | 0.03 | 0.63 | 0.01 |
| rs7181610 | 0.15 | 0.03 | 0.74 | 0.02 |
| rs8567 | 0.15 | 0.03 | 0.87 | 0.03 |
| rs13047416 | 0.18 | 0.06 | 0.52 | 0.002 |
| rs12602912 | 0.19 | 0.06 | 0.58 | 0.003 |
| rs2781668 | 0.19 | 0.06 | 0.68 | 0.01 |
| rs998732 | 0.19 | 0.05 | 0.69 | 0.01 |
| rs16906845 | 0.20 | 0.05 | 0.89 | 0.03 |
| rs2283093 | 0.20 | 0.04 | 0.92 | 0.04 |
| rs273697 | 0.20 | 0.04 | 1.01 | 0.05 |
| rs3851083 | 0.20 | 0.04 | 0.93 | 0.04 |
| rs6764533 | 0.20 | 0.05 | 0.85 | 0.03 |
| rs10838122 | 0.21 | 0.05 | 0.94 | 0.04 |
| rs12609744 | 0.21 | 0.05 | 0.84 | 0.03 |
| rs9463175 | 0.22 | 0.05 | 1.02 | 0.05 |
| rs881301 | 0.23 | 0.04 | 1.22 | 0.08 |
| rs9299 | 0.23 | 0.06 | 0.89 | 0.03 |
| rs1000096 | 0.24 | 0.08 | 0.75 | 0.01 |
| rs12776880 | 0.24 | 0.06 | 0.96 | 0.04 |
| rs6879326 | 0.25 | 0.05 | 1.31 | 0.10 |
| rs6138482 | 0.26 | 0.06 | 1.03 | 0.06 |
| rs903959 | 0.26 | 0.06 | 1.23 | 0.09 |
| rs12694021 | 0.27 | 0.06 | 1.27 | 0.10 |
| rs1862451 | 0.27 | 0.07 | 1.02 | 0.05 |
| rs2543132 | 0.27 | 0.07 | 1.06 | 0.06 |
| rs3915844 | 0.27 | 0.06 | 1.26 | 0.10 |
| rs4670626 | 0.27 | 0.06 | 1.26 | 0.10 |
| rs1006353 | 0.28 | 0.07 | 1.17 | 0.08 |
| rs7973955 | 0.28 | 0.07 | 1.17 | 0.08 |
| rs262956 | 0.29 | 0.08 | 1.10 | 0.07 |
| rs7083450 | 0.29 | 0.07 | 1.13 | 0.07 |
| rs7715256 | 0.30 | 0.12 | 0.79 | 0.01 |
| rs10886017 | 0.31 | 0.09 | 1.06 | 0.06 |
| rs12044597 | 0.31 | 0.09 | 1.08 | 0.07 |
| rs6587552 | 0.31 | 0.10 | 0.90 | 0.03 |
| rs7133378 | 0.31 | 0.08 | 1.26 | 0.10 |
| rs10118701 | 0.32 | 0.11 | 0.89 | 0.03 |
| rs11866815 | 0.32 | 0.10 | 1.04 | 0.06 |
| rs40245 | 0.32 | 0.07 | 1.46 | 0.14 |
| rs7710595 | 0.32 | 0.07 | 1.59 | 0.16 |
| rs8097544 | 0.32 | 0.10 | 0.96 | 0.04 |
| rs17757975 | 0.33 | 0.07 | 1.66 | 0.18 |
| rs1951455 | 0.33 | 0.10 | 1.12 | 0.08 |
| rs3814883 | 0.33 | 0.16 | 0.67 | 0.002 |
| rs1899689 | 0.34 | 0.08 | 1.43 | 0.14 |
| rs419261 | 0.34 | 0.08 | 1.53 | 0.16 |
| rs1040881 | 0.35 | 0.06 | 1.92 | 0.23 |
| rs6921533 | 0.35 | 0.08 | 1.51 | 0.16 |
| rs4372836 | 0.36 | 0.11 | 1.18 | 0.09 |
| rs7560871 | 0.36 | 0.09 | 1.46 | 0.15 |
| rs13110266 | 0.37 | 0.10 | 1.44 | 0.15 |
| rs10915840 | 0.38 | 0.08 | 1.74 | 0.21 |
| rs10953620 | 0.38 | 0.08 | 1.83 | 0.23 |
| rs11600990 | 0.38 | 0.11 | 1.28 | 0.12 |
| rs13292976 | 0.38 | 0.11 | 1.30 | 0.12 |
| rs1814170 | 0.38 | 0.10 | 1.42 | 0.15 |
| rs1866510 | 0.38 | 0.08 | 1.83 | 0.23 |
| rs450231 | 0.38 | 0.09 | 1.59 | 0.19 |
| rs719802 | 0.38 | 0.08 | 1.92 | 0.24 |
| rs7683836 | 0.38 | 0.09 | 1.66 | 0.20 |
| rs10744146 | 0.39 | 0.11 | 1.40 | 0.15 |
| rs6477694 | 0.39 | 0.11 | 1.44 | 0.16 |
| rs7124681 | 0.39 | 0.21 | 0.73 | 0.003 |
| rs13263601 | 0.40 | 0.14 | 1.17 | 0.09 |
| rs483752 | 0.40 | 0.08 | 2.04 | 0.27 |
| rs10460960 | 0.41 | 0.10 | 1.61 | 0.20 |
| rs11577179 | 0.41 | 0.09 | 1.84 | 0.24 |
| rs17820822 | 0.41 | 0.13 | 1.3 | 0.13 |
| rs2030342 | 0.41 | 0.13 | 1.29 | 0.13 |
| rs10131890 | 0.42 | 0.10 | 1.80 | 0.24 |
| rs10779751 | 0.42 | 0.11 | 1.53 | 0.19 |
| rs10935143 | 0.42 | 0.09 | 1.98 | 0.27 |
| rs1625427 | 0.42 | 0.12 | 1.50 | 0.18 |
| rs1956151 | 0.42 | 0.09 | 1.97 | 0.27 |
| rs2170382 | 0.42 | 0.09 | 1.87 | 0.25 |
| rs4864201 | 0.42 | 0.13 | 1.33 | 0.14 |
| rs6442101 | 0.42 | 0.10 | 1.83 | 0.25 |
| rs9426003 | 0.43 | 0.08 | 2.18 | 0.31 |
| rs10438964 | 0.44 | 0.11 | 1.74 | 0.24 |
| rs10754210 | 0.44 | 0.11 | 1.72 | 0.24 |
| rs12443621 | 0.44 | 0.08 | 2.32 | 0.34 |
| rs4936671 | 0.44 | 0.09 | 2.11 | 0.31 |
| rs1218822 | 0.45 | 0.16 | 1.23 | 0.12 |
| rs17538472 | 0.45 | 0.10 | 2.03 | 0.30 |
| rs2190788 | 0.46 | 0.14 | 1.54 | 0.21 |
| rs4796243 | 0.46 | 0.11 | 1.96 | 0.29 |
| rs12939549 | 0.47 | 0.19 | 1.17 | 0.10 |
| rs17207196 | 0.47 | 0.22 | 0.96 | 0.04 |
| rs3902951 | 0.47 | 0.12 | 1.82 | 0.27 |
| rs6738445 | 0.47 | 0.13 | 1.75 | 0.26 |
| rs1048932 | 0.48 | 0.18 | 1.31 | 0.15 |
| rs6700838 | 0.48 | 0.19 | 1.23 | 0.13 |
| rs6767619 | 0.48 | 0.12 | 1.94 | 0.30 |
| rs785278 | 0.48 | 0.13 | 1.75 | 0.27 |
| rs7893571 | 0.48 | 0.13 | 1.69 | 0.25 |
| rs9571687 | 0.48 | 0.13 | 1.80 | 0.27 |
| rs7042372 | 0.49 | 0.13 | 1.92 | 0.31 |
| rs7444298 | 0.49 | 0.18 | 1.35 | 0.17 |
| rs331949 | 0.50 | 0.11 | 2.26 | 0.36 |
| rs11496125 | 0.50 | 0.19 | 1.31 | 0.16 |
| rs12189178 | 0.50 | 0.16 | 1.60 | 0.24 |
| rs10797115 | 0.51 | 0.14 | 1.83 | 0.30 |
| rs3810291 | 0.51 | 0.28 | 0.95 | 0.03 |
| rs7209235 | 0.51 | 0.09 | 2.85 | 0.45 |
| rs7869771 | 0.51 | 0.14 | 1.88 | 0.31 |
| rs11792069 | 0.52 | 0.11 | 2.56 | 0.42 |
| rs761423 | 0.52 | 0.12 | 2.19 | 0.37 |
| rs9630985 | 0.52 | 0.20 | 1.35 | 0.18 |
| rs17094222 | 0.53 | 0.18 | 1.58 | 0.25 |
| rs6968554 | 0.53 | 0.10 | 2.71 | 0.45 |
| rs3134438 | 0.54 | 0.10 | 2.8 | 0.46 |
| rs11781222 | 0.55 | 0.12 | 2.46 | 0.43 |
| rs11790280 | 0.55 | 0.11 | 2.81 | 0.47 |
| rs7181498 | 0.55 | 0.20 | 1.50 | 0.24 |
| rs8036040 | 0.55 | 0.12 | 2.59 | 0.45 |
| rs972283 | 0.55 | 0.10 | 3.20 | 0.51 |
| rs10920678 | 0.56 | 0.20 | 1.61 | 0.28 |
| rs6577584 | 0.56 | 0.15 | 2.04 | 0.38 |
| rs7903146 | 0.56 | 0.21 | 1.48 | 0.24 |
| rs10929925 | 0.57 | 0.18 | 1.78 | 0.34 |
| rs12041258 | 0.57 | 0.16 | 2.06 | 0.39 |
| rs12446632 | 0.57 | 0.29 | 1.11 | 0.10 |
| rs17531363 | 0.57 | 0.16 | 2.07 | 0.40 |
| rs4676084 | 0.57 | 0.11 | 2.94 | 0.50 |
| rs4820408 | 0.57 | 0.20 | 1.64 | 0.30 |
| rs1000940 | 0.58 | 0.19 | 1.74 | 0.33 |
| rs13021737 | 0.58 | 0.41 | 0.84 | 0.004 |
| rs6474945 | 0.58 | 0.25 | 1.34 | 0.20 |
| rs7024334 | 0.58 | 0.15 | 2.22 | 0.42 |
| rs9816226 | 0.58 | 0.30 | 1.10 | 0.10 |
| rs17069831 | 0.59 | 0.10 | 3.36 | 0.55 |
| rs17724992 | 0.59 | 0.22 | 1.57 | 0.29 |
| rs17806379 | 0.59 | 0.26 | 1.35 | 0.21 |
| rs1789165 | 0.59 | 0.18 | 1.94 | 0.38 |
| rs2275426 | 0.59 | 0.13 | 2.66 | 0.49 |
| rs4722398 | 0.59 | 0.13 | 2.72 | 0.50 |
| rs4722672 | 0.59 | 0.15 | 2.30 | 0.45 |
| rs6019482 | 0.59 | 0.17 | 2.00 | 0.40 |
| rs3781099 | 0.60 | 0.15 | 2.37 | 0.47 |
| rs4886869 | 0.60 | 0.12 | 3.00 | 0.53 |
| rs577525 | 0.60 | 0.22 | 1.63 | 0.31 |
| rs6121381 | 0.60 | 0.12 | 2.91 | 0.52 |
| rs10883759 | 0.61 | 0.15 | 2.45 | 0.48 |
| rs12655756 | 0.61 | 0.17 | 2.18 | 0.44 |
| rs1451533 | 0.61 | 0.20 | 1.83 | 0.38 |
| rs2192158 | 0.61 | 0.17 | 2.18 | 0.45 |
| rs4764949 | 0.61 | 0.25 | 1.49 | 0.28 |
| rs930295 | 0.61 | 0.22 | 1.69 | 0.34 |
| rs10132280 | 0.62 | 0.28 | 1.35 | 0.22 |
| rs10275044 | 0.62 | 0.12 | 3.12 | 0.56 |
| rs11736228 | 0.62 | 0.17 | 2.28 | 0.47 |
| rs1884389 | 0.62 | 0.13 | 2.87 | 0.54 |
| rs4518345 | 0.62 | 0.13 | 2.97 | 0.55 |
| rs7006629 | 0.62 | 0.14 | 2.66 | 0.52 |
| rs7123876 | 0.62 | 0.14 | 2.89 | 0.55 |
| rs7206608 | 0.62 | 0.17 | 2.22 | 0.46 |
| rs9349239 | 0.62 | 0.17 | 2.32 | 0.48 |
| rs12042959 | 0.63 | 0.13 | 2.92 | 0.55 |
| rs12429545 | 0.63 | 0.29 | 1.37 | 0.25 |
| rs2605603 | 0.63 | 0.14 | 2.88 | 0.55 |
| rs3807566 | 0.63 | 0.18 | 2.28 | 0.48 |
| rs4372296 | 0.63 | 0.15 | 2.54 | 0.51 |
| rs13329567 | 0.64 | 0.33 | 1.24 | 0.19 |
| rs2842385 | 0.64 | 0.12 | 3.35 | 0.60 |
| rs3803286 | 0.64 | 0.25 | 1.60 | 0.34 |
| rs7607351 | 0.64 | 0.16 | 2.45 | 0.51 |
| rs10883553 | 0.65 | 0.17 | 2.45 | 0.52 |
| rs16932761 | 0.65 | 0.17 | 2.41 | 0.52 |
| rs10818810 | 0.66 | 0.17 | 2.56 | 0.55 |
| rs17014375 | 0.66 | 0.17 | 2.60 | 0.55 |
| rs17035438 | 0.66 | 0.16 | 2.71 | 0.56 |
| rs2122042 | 0.66 | 0.29 | 1.53 | 0.34 |
| rs248139 | 0.66 | 0.14 | 3.11 | 0.60 |
| rs8016859 | 0.66 | 0.19 | 2.30 | 0.52 |
| rs11079849 | 0.67 | 0.27 | 1.70 | 0.40 |
| rs4936175 | 0.67 | 0.18 | 2.48 | 0.55 |
| rs6827083 | 0.67 | 0.12 | 3.73 | 0.65 |
| rs10989568 | 0.68 | 0.16 | 2.92 | 0.60 |
| rs1941697 | 0.68 | 0.19 | 2.42 | 0.55 |
| rs3849570 | 0.68 | 0.19 | 2.40 | 0.55 |
| rs4759073 | 0.68 | 0.17 | 2.69 | 0.59 |
| rs555267 | 0.68 | 0.18 | 2.56 | 0.57 |
| rs733594 | 0.68 | 0.18 | 2.59 | 0.57 |
| rs40067 | 0.69 | 0.32 | 1.50 | 0.35 |
| rs11150911 | 0.70 | 0.19 | 2.57 | 0.59 |
| rs4740383 | 0.71 | 0.19 | 2.67 | 0.61 |
| rs6804181 | 0.71 | 0.18 | 2.75 | 0.62 |
| rs10832778 | 0.72 | 0.20 | 2.65 | 0.62 |
| rs10923724 | 0.72 | 0.19 | 2.78 | 0.64 |
| rs1320903 | 0.72 | 0.32 | 1.64 | 0.44 |
| rs9927848 | 0.72 | 0.16 | 3.31 | 0.67 |
| rs1431659 | 0.73 | 0.28 | 1.87 | 0.51 |
| rs663129 | 0.73 | 0.52 | 1.03 | 0.07 |
| rs902695 | 0.73 | 0.16 | 3.42 | 0.69 |
| rs4303732 | 0.74 | 0.23 | 2.39 | 0.62 |
| rs12042908 | 0.75 | 0.32 | 1.79 | 0.52 |
| rs12885454 | 0.75 | 0.31 | 1.84 | 0.54 |
| rs4906908 | 0.75 | 0.16 | 3.52 | 0.72 |
| rs11753081 | 0.76 | 0.17 | 3.35 | 0.72 |
| rs2307111 | 0.76 | 0.41 | 1.39 | 0.37 |
| rs12675063 | 0.77 | 0.15 | 3.99 | 0.75 |
| rs1296328 | 0.77 | 0.31 | 1.89 | 0.57 |
| rs13107325 | 0.77 | 0.40 | 1.49 | 0.44 |
| rs329277 | 0.77 | 0.16 | 3.75 | 0.75 |
| rs3800649 | 0.77 | 0.16 | 3.77 | 0.75 |
| rs7760082 | 0.77 | 0.19 | 3.06 | 0.71 |
| rs2820311 | 0.79 | 0.39 | 1.62 | 0.52 |
| rs6804842 | 0.79 | 0.27 | 2.34 | 0.68 |
| rs8047395 | 0.79 | 0.62 | 1.01 | 0.06 |
| rs8089514 | 0.79 | 0.20 | 3.12 | 0.74 |
| rs9538141 | 0.79 | 0.29 | 2.18 | 0.65 |
| rs13227658 | 0.80 | 0.28 | 2.24 | 0.66 |
| rs4814512 | 0.80 | 0.18 | 3.59 | 0.77 |
| rs11987383 | 0.81 | 0.16 | 4.23 | 0.80 |
| rs13209872 | 0.81 | 0.27 | 2.42 | 0.71 |
| rs4673553 | 0.81 | 0.26 | 2.51 | 0.71 |
| rs7865157 | 0.81 | 0.17 | 3.93 | 0.79 |
| rs7941030 | 0.81 | 0.18 | 3.57 | 0.78 |
| rs1973993 | 0.82 | 0.37 | 1.80 | 0.62 |
| rs3736485 | 0.82 | 0.25 | 2.69 | 0.75 |
| rs6690764 | 0.82 | 0.24 | 2.83 | 0.76 |
| rs13290794 | 0.83 | 0.25 | 2.69 | 0.75 |
| rs2850969 | 0.83 | 0.21 | 3.27 | 0.79 |
| rs1455137 | 0.84 | 0.18 | 3.88 | 0.82 |
| rs156151 | 0.84 | 0.24 | 2.95 | 0.78 |
| rs2890652 | 0.84 | 0.25 | 2.90 | 0.79 |
| rs13174863 | 0.85 | 0.24 | 3.02 | 0.80 |
| rs1668633 | 0.85 | 0.16 | 4.42 | 0.85 |
| rs2304130 | 0.85 | 0.16 | 4.58 | 0.85 |
| rs7084454 | 0.85 | 0.36 | 2.02 | 0.72 |
| rs7600699 | 0.85 | 0.19 | 3.69 | 0.82 |
| rs9367368 | 0.85 | 0.21 | 3.47 | 0.82 |
| rs1362910 | 0.86 | 0.22 | 3.31 | 0.83 |
| rs9688431 | 0.86 | 0.20 | 3.61 | 0.84 |
| rs12587412 | 0.87 | 0.29 | 2.57 | 0.80 |
| rs977540 | 0.87 | 0.23 | 3.32 | 0.83 |
| rs208015 | 0.89 | 0.39 | 2.01 | 0.77 |
| rs7871866 | 0.89 | 0.27 | 2.97 | 0.85 |
| rs11915371 | 0.90 | 0.25 | 3.26 | 0.87 |
| rs1323068 | 0.90 | 0.20 | 4.02 | 0.89 |
| rs2357760 | 0.90 | 0.28 | 2.83 | 0.85 |
| rs7557796 | 0.90 | 0.32 | 2.55 | 0.84 |
| rs987237 | 0.90 | 0.55 | 1.46 | 0.66 |
| rs2733287 | 0.91 | 0.34 | 2.44 | 0.85 |
| rs6786582 | 0.91 | 0.30 | 2.76 | 0.87 |
| rs3753549 | 0.92 | 0.30 | 2.83 | 0.89 |
| rs6877851 | 0.92 | 0.21 | 4.00 | 0.91 |
| rs11945861 | 0.93 | 0.27 | 3.29 | 0.92 |
| rs2010281 | 0.93 | 0.32 | 2.68 | 0.89 |
| rs6720868 | 0.93 | 0.31 | 2.76 | 0.90 |
| rs11635675 | 0.94 | 0.24 | 3.59 | 0.93 |
| rs2731222 | 0.94 | 0.22 | 3.96 | 0.93 |
| rs6448587 | 0.95 | 0.29 | 3.12 | 0.94 |
| rs7195386 | 0.95 | 0.26 | 3.42 | 0.94 |
| rs1405348 | 0.96 | 0.45 | 2.07 | 0.92 |
| rs1658820 | 0.96 | 0.25 | 3.64 | 0.95 |
| rs11118308 | 0.97 | 0.20 | 4.77 | 0.97 |
| rs1608445 | 0.97 | 0.19 | 5.10 | 0.97 |
| rs2838006 | 0.97 | 0.27 | 3.52 | 0.96 |
| rs7531656 | 0.97 | 0.40 | 2.32 | 0.95 |
| rs895330 | 0.97 | 0.34 | 2.74 | 0.95 |
| rs925421 | 0.97 | 0.19 | 4.89 | 0.97 |
| rs1700137 | 0.98 | 0.24 | 4.09 | 0.98 |
| rs2304607 | 0.98 | 0.48 | 2.01 | 0.96 |
| rs6265 | 0.99 | 0.61 | 1.60 | 0.96 |
| rs7334078 | 0.99 | 0.23 | 4.26 | 0.99 |
| rs8071182 | 0.99 | 0.19 | 5.17 | 0.99 |
| rs8081039 | 0.99 | 0.24 | 4.02 | 0.99 |
| rs2479958 | 1.00 | 0.35 | 2.88 | 1.00 |
| rs7187776 | 1.00 | 0.54 | 1.85 | 1.00 |
| rs7805441 | 1.00 | 0.24 | 4.21 | 1.00 |
| rs12514473 | 1.00 | 0.32 | 3.12 | 1.00 |
| rs17681708 | 1.01 | 0.19 | 5.33 | 0.99 |
| rs7607369 | 1.01 | 0.25 | 4.05 | 0.99 |
| rs8123881 | 1.01 | 0.29 | 3.49 | 0.99 |
| rs889398 | 1.01 | 0.44 | 2.31 | 0.99 |
| rs901630 | 1.01 | 0.33 | 3.09 | 0.98 |
| rs9077 | 1.01 | 0.28 | 3.64 | 0.98 |
| rs10938397 | 1.02 | 0.62 | 1.66 | 0.94 |
| rs10268050 | 1.03 | 0.18 | 6.02 | 0.98 |
| rs12448738 | 1.03 | 0.26 | 4.03 | 0.97 |
| rs1358808 | 1.03 | 0.32 | 3.28 | 0.96 |
| rs6713781 | 1.03 | 0.28 | 3.82 | 0.96 |
| rs10408013 | 1.04 | 0.21 | 5.17 | 0.96 |
| rs4858193 | 1.04 | 0.25 | 4.27 | 0.96 |
| rs6443750 | 1.04 | 0.21 | 5.17 | 0.96 |
| rs6843738 | 1.04 | 0.22 | 4.84 | 0.96 |
| rs6919443 | 1.04 | 0.19 | 5.79 | 0.96 |
| rs9530843 | 1.04 | 0.28 | 3.88 | 0.95 |
| rs1477887 | 1.05 | 0.33 | 3.32 | 0.93 |
| rs1657930 | 1.06 | 0.22 | 5.21 | 0.94 |
| rs1884897 | 1.06 | 0.45 | 2.51 | 0.89 |
| rs9931967 | 1.06 | 0.40 | 2.85 | 0.90 |
| rs998584 | 1.07 | 0.32 | 3.60 | 0.91 |
| rs10878946 | 1.08 | 0.30 | 3.94 | 0.91 |
| rs6548221 | 1.08 | 0.31 | 3.79 | 0.91 |
| rs7144011 | 1.08 | 0.55 | 2.14 | 0.82 |
| rs2154297 | 1.09 | 0.21 | 5.60 | 0.92 |
| rs9294260 | 1.09 | 0.37 | 3.20 | 0.88 |
| rs7694732 | 1.10 | 0.22 | 5.44 | 0.91 |
| rs13240600 | 1.12 | 0.40 | 3.16 | 0.83 |
| rs711347 | 1.12 | 0.26 | 4.92 | 0.88 |
| rs1412235 | 1.14 | 0.57 | 2.29 | 0.70 |
| rs9951893 | 1.14 | 0.29 | 4.53 | 0.85 |
| rs10146527 | 1.15 | 0.35 | 3.82 | 0.82 |
| rs2836961 | 1.15 | 0.24 | 5.55 | 0.86 |
| rs6710871 | 1.15 | 0.31 | 4.23 | 0.83 |
| rs16851483 | 1.16 | 0.49 | 2.77 | 0.74 |
| rs7948120 | 1.16 | 0.29 | 4.70 | 0.83 |
| rs1365466 | 1.19 | 0.31 | 4.51 | 0.80 |
| rs427943 | 1.20 | 0.47 | 3.05 | 0.70 |
| rs4969387 | 1.20 | 0.34 | 4.15 | 0.78 |
| rs9806058 | 1.20 | 0.28 | 5.16 | 0.81 |
| rs11074446 | 1.21 | 0.43 | 3.38 | 0.72 |
| rs1853639 | 1.21 | 0.27 | 5.33 | 0.80 |
| rs12364470 | 1.22 | 0.32 | 4.56 | 0.77 |
| rs12595158 | 1.22 | 0.38 | 3.88 | 0.74 |
| rs6888159 | 1.23 | 0.26 | 5.8 | 0.79 |
| rs1522569 | 1.24 | 0.32 | 4.78 | 0.76 |
| rs7147503 | 1.25 | 0.32 | 4.92 | 0.74 |
| rs7899106 | 1.25 | 0.40 | 3.86 | 0.70 |
| rs827092 | 1.25 | 0.37 | 4.29 | 0.72 |
| rs1437929 | 1.26 | 0.34 | 4.72 | 0.73 |
| rs7217226 | 1.26 | 0.35 | 4.54 | 0.72 |
| rs3739733 | 1.29 | 0.32 | 5.30 | 0.72 |
| rs7844647 | 1.29 | 0.30 | 5.57 | 0.74 |
| rs10182181 | 1.31 | 0.79 | 2.17 | 0.30 |
| rs775731 | 1.31 | 0.27 | 6.34 | 0.74 |
| rs11672660 | 1.32 | 0.73 | 2.38 | 0.35 |
| rs1730859 | 1.32 | 0.32 | 5.52 | 0.70 |
| rs326889 | 1.32 | 0.38 | 4.59 | 0.66 |
| rs4718966 | 1.32 | 0.37 | 4.67 | 0.67 |
| rs6564360 | 1.32 | 0.27 | 6.40 | 0.73 |
| rs925018 | 1.33 | 0.36 | 4.86 | 0.67 |
| rs1106908 | 1.34 | 0.44 | 4.09 | 0.61 |
| rs28350 | 1.34 | 0.38 | 4.69 | 0.65 |
| rs6700816 | 1.34 | 0.32 | 5.50 | 0.69 |
| rs7599312 | 1.34 | 0.51 | 3.49 | 0.55 |
| rs11611496 | 1.35 | 0.41 | 4.47 | 0.62 |
| rs4639527 | 1.35 | 0.47 | 3.86 | 0.57 |
| rs543874 | 1.35 | 0.89 | 2.05 | 0.15 |
| rs12577642 | 1.36 | 0.61 | 3.04 | 0.45 |
| rs1782507 | 1.36 | 0.40 | 4.62 | 0.62 |
| rs10962549 | 1.37 | 0.44 | 4.23 | 0.59 |
| rs13247665 | 1.37 | 0.41 | 4.55 | 0.60 |
| rs4366093 | 1.37 | 0.34 | 5.51 | 0.66 |
| rs7784465 | 1.37 | 0.34 | 5.63 | 0.66 |
| rs11668301 | 1.38 | 0.48 | 3.97 | 0.55 |
| rs2033529 | 1.38 | 0.55 | 3.45 | 0.49 |
| rs2367112 | 1.38 | 0.36 | 5.22 | 0.64 |
| rs29938 | 1.38 | 0.45 | 4.24 | 0.58 |
| rs12731372 | 1.41 | 0.28 | 7.15 | 0.68 |
| rs17446091 | 1.41 | 0.28 | 7.03 | 0.68 |
| rs1045411 | 1.42 | 0.37 | 5.49 | 0.61 |
| rs6461115 | 1.42 | 0.39 | 5.19 | 0.59 |
| rs10211055 | 1.43 | 0.46 | 4.45 | 0.54 |
| rs2143253 | 1.43 | 0.40 | 5.04 | 0.58 |
| rs215632 | 1.45 | 0.49 | 4.35 | 0.50 |
| rs6545714 | 1.45 | 0.62 | 3.40 | 0.39 |
| rs6711584 | 1.45 | 0.43 | 4.95 | 0.55 |
| rs3822683 | 1.46 | 0.38 | 5.65 | 0.58 |
| rs7531118 | 1.47 | 0.78 | 2.78 | 0.23 |
| rs762147 | 1.47 | 0.25 | 8.63 | 0.67 |
| rs9603697 | 1.47 | 0.44 | 4.92 | 0.53 |
| rs7615297 | 1.48 | 0.33 | 6.61 | 0.61 |
| rs9332817 | 1.48 | 0.35 | 6.31 | 0.60 |
| rs2429150 | 1.49 | 0.33 | 6.79 | 0.61 |
| rs2124499 | 1.50 | 0.37 | 6.10 | 0.57 |
| rs4516268 | 1.51 | 0.59 | 3.86 | 0.39 |
| rs2481665 | 1.52 | 0.57 | 4.06 | 0.41 |
| rs7138803 | 1.52 | 0.89 | 2.60 | 0.12 |
| rs1452075 | 1.53 | 0.43 | 5.50 | 0.51 |
| rs4653017 | 1.53 | 0.38 | 6.20 | 0.55 |
| rs7117238 | 1.53 | 0.31 | 7.49 | 0.60 |
| rs900144 | 1.53 | 0.59 | 3.99 | 0.38 |
| rs9507983 | 1.53 | 0.53 | 4.39 | 0.43 |
| rs6235 | 1.54 | 0.54 | 4.40 | 0.42 |
| rs12564992 | 1.55 | 0.43 | 5.58 | 0.50 |
| rs13191362 | 1.55 | 0.54 | 4.46 | 0.41 |
| rs2228213 | 1.55 | 0.47 | 5.07 | 0.47 |
| rs2365389 | 1.55 | 0.61 | 3.9 | 0.35 |
| rs7975187 | 1.55 | 0.34 | 7.05 | 0.57 |
| rs12147845 | 1.56 | 0.45 | 5.46 | 0.48 |
| rs4744275 | 1.59 | 0.44 | 5.72 | 0.48 |
| rs7550711 | 1.59 | 0.78 | 3.23 | 0.20 |
| rs9362662 | 1.59 | 0.38 | 6.68 | 0.53 |
| rs1512065 | 1.60 | 0.36 | 7.12 | 0.54 |
| rs1982441 | 1.60 | 0.44 | 5.79 | 0.48 |
| rs1852006 | 1.61 | 0.55 | 4.68 | 0.38 |
| rs13072095 | 1.62 | 0.25 | 10.61 | 0.61 |
| rs934515 | 1.66 | 0.45 | 6.12 | 0.44 |
| rs9595908 | 1.66 | 0.60 | 4.63 | 0.33 |
| rs1075901 | 1.67 | 0.45 | 6.20 | 0.44 |
| rs159032 | 1.67 | 0.40 | 6.96 | 0.48 |
| rs1928295 | 1.67 | 0.54 | 5.14 | 0.37 |
| rs12420725 | 1.68 | 0.39 | 7.27 | 0.49 |
| rs12575252 | 1.68 | 0.68 | 4.14 | 0.26 |
| rs12033257 | 1.72 | 0.53 | 5.52 | 0.36 |
| rs845084 | 1.72 | 0.47 | 6.33 | 0.41 |
| rs453520 | 1.73 | 0.55 | 5.38 | 0.35 |
| rs12101393 | 1.74 | 0.37 | 8.12 | 0.48 |
| rs11855853 | 1.76 | 0.50 | 6.19 | 0.38 |
| rs573455 | 1.77 | 0.30 | 10.38 | 0.53 |
| rs740157 | 1.77 | 0.46 | 6.83 | 0.41 |
| rs9291467 | 1.77 | 0.57 | 5.50 | 0.32 |
| rs10467530 | 1.80 | 0.33 | 9.81 | 0.50 |
| rs12035149 | 1.81 | 0.41 | 8.05 | 0.43 |
| rs7211567 | 1.81 | 0.47 | 6.99 | 0.39 |
| rs967605 | 1.81 | 0.57 | 5.74 | 0.31 |
| rs2012927 | 1.82 | 0.51 | 6.51 | 0.36 |
| rs498240 | 1.83 | 0.56 | 6.00 | 0.32 |
| rs9475173 | 1.84 | 0.39 | 8.62 | 0.44 |
| rs12680842 | 1.87 | 0.53 | 6.63 | 0.33 |
| rs7239114 | 1.89 | 0.48 | 7.48 | 0.36 |
| rs16966801 | 1.91 | 0.52 | 6.97 | 0.33 |
| rs954018 | 1.91 | 0.51 | 7.08 | 0.33 |
| rs1003081 | 1.94 | 0.51 | 7.31 | 0.33 |
| rs7640424 | 1.94 | 0.56 | 6.69 | 0.30 |
| rs217671 | 1.95 | 0.51 | 7.39 | 0.33 |
| rs7164727 | 1.95 | 0.77 | 4.99 | 0.16 |
| rs223051 | 1.97 | 0.43 | 9.03 | 0.38 |
| rs9544915 | 1.97 | 0.36 | 10.72 | 0.43 |
| rs340025 | 1.98 | 0.53 | 7.49 | 0.31 |
| rs6449531 | 1.98 | 0.53 | 7.37 | 0.31 |
| rs8102137 | 1.99 | 0.84 | 4.72 | 0.12 |
| rs4671358 | 2.00 | 0.40 | 10.01 | 0.40 |
| rs2619976 | 2.02 | 0.38 | 10.60 | 0.41 |
| rs7601895 | 2.02 | 0.63 | 6.52 | 0.24 |
| rs1371108 | 2.06 | 0.50 | 8.49 | 0.32 |
| rs2230590 | 2.06 | 1.07 | 3.97 | 0.03 |
| rs4725984 | 2.10 | 0.57 | 7.72 | 0.26 |
| rs3800229 | 2.13 | 0.78 | 5.83 | 0.14 |
| rs17327461 | 2.16 | 0.61 | 7.68 | 0.24 |
| rs284227 | 2.16 | 0.64 | 7.26 | 0.21 |
| rs1241986 | 2.17 | 0.44 | 10.70 | 0.34 |
| rs4307239 | 2.17 | 0.54 | 8.77 | 0.28 |
| rs10930641 | 2.18 | 0.66 | 7.26 | 0.20 |
| rs10083803 | 2.25 | 0.54 | 9.31 | 0.26 |
| rs1229057 | 2.25 | 0.53 | 9.63 | 0.27 |
| rs4986044 | 2.25 | 0.83 | 6.07 | 0.11 |
| rs1544459 | 2.28 | 0.46 | 11.29 | 0.31 |
| rs1937684 | 2.29 | 0.51 | 10.33 | 0.28 |
| rs7711753 | 2.29 | 0.70 | 7.49 | 0.17 |
| rs7730898 | 2.29 | 0.80 | 6.54 | 0.12 |
| rs12098284 | 2.38 | 0.59 | 9.62 | 0.23 |
| rs2080454 | 2.40 | 0.68 | 8.47 | 0.17 |
| rs7779498 | 2.41 | 0.40 | 14.39 | 0.33 |
| rs1014194 | 2.51 | 0.48 | 13.12 | 0.27 |
| rs9814633 | 2.52 | 0.61 | 10.38 | 0.20 |
| rs1899951 | 2.55 | 0.65 | 10.05 | 0.18 |
| rs7874154 | 2.57 | 0.70 | 9.41 | 0.15 |
| rs4148155 | 2.63 | 0.69 | 10.10 | 0.16 |
| rs12922346 | 2.68 | 0.65 | 11.00 | 0.17 |
| rs4624596 | 2.70 | 0.63 | 11.57 | 0.18 |
| rs12439798 | 2.72 | 0.76 | 9.68 | 0.12 |
| rs12964689 | 2.72 | 1.24 | 5.94 | 0.01 |
| rs11251352 | 2.77 | 0.59 | 13.00 | 0.20 |
| rs273504 | 2.77 | 0.98 | 7.82 | 0.05 |
| rs11889536 | 2.78 | 0.81 | 9.54 | 0.10 |
| rs11783247 | 2.83 | 1.09 | 7.33 | 0.03 |
| rs17681451 | 2.83 | 0.63 | 12.72 | 0.17 |
| rs7102454 | 2.88 | 1.00 | 8.26 | 0.05 |
| rs1927790 | 2.89 | 0.95 | 8.79 | 0.06 |
| rs10830452 | 2.92 | 0.63 | 13.58 | 0.17 |
| rs12593036 | 2.94 | 0.95 | 9.12 | 0.06 |
| rs972540 | 2.94 | 0.71 | 12.11 | 0.14 |
| rs1038088 | 2.99 | 0.77 | 11.60 | 0.11 |
| rs7674623 | 3.04 | 0.70 | 13.16 | 0.14 |
| rs6900723 | 3.08 | 0.67 | 14.12 | 0.15 |
| rs7535528 | 3.08 | 0.99 | 9.58 | 0.05 |
| rs11075489 | 3.11 | 0.74 | 13.01 | 0.12 |
| rs11659764 | 3.20 | 0.74 | 13.82 | 0.12 |
| rs2721965 | 3.34 | 1.27 | 8.74 | 0.01 |
| rs4766710 | 3.35 | 0.74 | 15.12 | 0.12 |
| rs1020548 | 3.39 | 0.67 | 17.08 | 0.14 |
| rs2100814 | 3.39 | 0.76 | 15.07 | 0.11 |
| rs849135 | 3.42 | 0.83 | 14.15 | 0.09 |
| rs13321566 | 3.50 | 0.72 | 17.02 | 0.12 |
| rs10867256 | 3.60 | 0.89 | 14.51 | 0.07 |
| rs25832 | 3.63 | 0.77 | 16.99 | 0.10 |
| rs1436343 | 3.66 | 1.16 | 11.61 | 0.03 |
| rs1035010 | 3.73 | 1.02 | 13.65 | 0.05 |
| rs12888545 | 3.84 | 0.99 | 14.88 | 0.05 |
| rs2425857 | 3.87 | 0.94 | 16.00 | 0.06 |
| rs2075650 | 4.06 | 1.57 | 10.48 | 0.004 |
| rs10269783 | 4.08 | 1.17 | 14.28 | 0.03 |
| rs1909586 | 4.12 | 0.81 | 20.95 | 0.09 |
| rs4985155 | 4.12 | 1.03 | 16.53 | 0.05 |
| rs756717 | 4.13 | 1.29 | 13.25 | 0.02 |
| rs3844598 | 4.19 | 0.79 | 22.26 | 0.09 |
| rs8070454 | 4.30 | 0.85 | 21.74 | 0.08 |
| rs8016771 | 4.32 | 1.01 | 18.50 | 0.05 |
| rs580438 | 4.33 | 0.94 | 19.99 | 0.06 |
| rs7556169 | 4.33 | 0.88 | 21.42 | 0.07 |
| rs11066301 | 4.37 | 1.05 | 18.19 | 0.04 |
| rs559267 | 4.54 | 0.95 | 21.78 | 0.06 |
| rs16828086 | 4.77 | 0.84 | 26.97 | 0.08 |
| rs10745785 | 4.84 | 1.08 | 21.70 | 0.04 |
| rs2875762 | 4.97 | 1.21 | 20.38 | 0.03 |
| rs11583122 | 5.11 | 1.06 | 24.64 | 0.04 |
| rs12448257 | 5.13 | 1.70 | 15.54 | 0.004 |
| rs17105272 | 5.14 | 1.07 | 24.64 | 0.04 |
| rs2282231 | 5.26 | 1.60 | 17.26 | 0.01 |
| rs11773362 | 5.85 | 1.3 | 26.22 | 0.02 |
| rs17551974 | 6.23 | 1.51 | 25.73 | 0.01 |
| rs2814992 | 6.25 | 3.19 | 12.24 | 9.03×10^-8^ |
| rs4284600 | 6.40 | 1.61 | 25.40 | 0.01 |
| rs10733051 | 7.02 | 1.31 | 37.55 | 0.02 |
| rs17203016 | 8.73 | 2.30 | 33.10 | 0.001 |
| rs2160077 | 10.09 | 1.87 | 54.48 | 0.01 |
| rs825680 | 10.35 | 1.93 | 55.36 | 0.01 |
| rs12615778 | 11.07 | 2.19 | 55.96 | 0.004 |
| rs10818938 | 14.14 | 3.57 | 55.96 | 1.6×10^-4^ |
| rs17636031 | 25.15 | 8.25 | 76.69 | 1.43×10^-8^ |

**Table S12: The MR association of individual SNPs of height with aggressive prostate cancer risk**

| **SNP** | **OR** | **LCI** | **UCI** | **P_assoc_** |
| --- | --- | --- | --- | --- |
| rs2763273 | 0.08 | 0.02 | 0.33 | 5.65×10^-4^ |
| rs12055154 | 0.10 | 0.02 | 0.44 | 0.003 |
| rs7899004 | 0.11 | 0.04 | 0.33 | 7.60×10^-5^ |
| rs12597498 | 0.15 | 0.03 | 0.72 | 0.02 |
| rs7033940 | 0.15 | 0.03 | 0.81 | 0.03 |
| rs1552173 | 0.19 | 0.04 | 0.94 | 0.04 |
| rs7727731 | 0.19 | 0.05 | 0.71 | 0.01 |
| rs1980850 | 0.20 | 0.06 | 0.69 | 0.01 |
| rs6688100 | 0.20 | 0.03 | 1.18 | 0.08 |
| rs7544462 | 0.22 | 0.05 | 1.03 | 0.05 |
| rs9309101 | 0.23 | 0.06 | 0.87 | 0.03 |
| rs11745439 | 0.24 | 0.07 | 0.78 | 0.02 |
| rs12639764 | 0.24 | 0.09 | 0.68 | 0.01 |
| rs761391 | 0.24 | 0.06 | 1.01 | 0.05 |
| rs960006 | 0.25 | 0.06 | 1.04 | 0.06 |
| rs7774834 | 0.27 | 0.06 | 1.23 | 0.09 |
| rs11642612 | 0.28 | 0.05 | 1.63 | 0.16 |
| rs2510396 | 0.28 | 0.07 | 1.14 | 0.07 |
| rs3132297 | 0.28 | 0.06 | 1.37 | 0.12 |
| rs12144094 | 0.29 | 0.06 | 1.31 | 0.11 |
| rs7112925 | 0.29 | 0.09 | 0.93 | 0.04 |
| rs2238300 | 0.30 | 0.08 | 1.11 | 0.07 |
| rs4344931 | 0.30 | 0.06 | 1.39 | 0.12 |
| rs4605213 | 0.30 | 0.06 | 1.44 | 0.13 |
| rs1614303 | 0.31 | 0.06 | 1.54 | 0.15 |
| rs3807931 | 0.31 | 0.11 | 0.85 | 0.02 |
| rs833152 | 0.32 | 0.06 | 1.73 | 0.18 |
| rs12987566 | 0.33 | 0.09 | 1.20 | 0.09 |
| rs4239020 | 0.33 | 0.07 | 1.49 | 0.15 |
| rs1550162 | 0.34 | 0.10 | 1.19 | 0.09 |
| rs10779751 | 0.35 | 0.08 | 1.56 | 0.17 |
| rs4803468 | 0.35 | 0.13 | 0.92 | 0.03 |
| rs1036477 | 0.36 | 0.08 | 1.58 | 0.18 |
| rs929637 | 0.36 | 0.08 | 1.69 | 0.20 |
| rs10495098 | 0.37 | 0.08 | 1.67 | 0.20 |
| rs12926008 | 0.38 | 0.08 | 1.77 | 0.22 |
| rs6733349 | 0.38 | 0.09 | 1.51 | 0.17 |
| rs10863936 | 0.39 | 0.10 | 1.56 | 0.18 |
| rs12779328 | 0.39 | 0.13 | 1.15 | 0.09 |
| rs6462432 | 0.39 | 0.07 | 2.08 | 0.27 |
| rs6902771 | 0.39 | 0.16 | 0.95 | 0.04 |
| rs780094 | 0.39 | 0.11 | 1.42 | 0.15 |
| rs11950938 | 0.40 | 0.10 | 1.60 | 0.20 |
| rs17113369 | 0.40 | 0.15 | 1.08 | 0.07 |
| rs749234 | 0.40 | 0.07 | 2.43 | 0.32 |
| rs9292468 | 0.41 | 0.18 | 0.93 | 0.03 |
| rs12344396 | 0.42 | 0.09 | 1.93 | 0.26 |
| rs2337143 | 0.42 | 0.08 | 2.16 | 0.30 |
| rs2149163 | 0.43 | 0.11 | 1.68 | 0.22 |
| rs4974480 | 0.43 | 0.14 | 1.29 | 0.13 |
| rs1055144 | 0.44 | 0.09 | 2.25 | 0.33 |
| rs4548838 | 0.44 | 0.19 | 1.00 | 0.05 |
| rs7261425 | 0.44 | 0.10 | 1.86 | 0.26 |
| rs7985356 | 0.44 | 0.11 | 1.75 | 0.24 |
| rs11783655 | 0.45 | 0.09 | 2.21 | 0.32 |
| rs7259684 | 0.45 | 0.10 | 1.95 | 0.29 |
| rs1461503 | 0.46 | 0.10 | 2.08 | 0.31 |
| rs4325879 | 0.46 | 0.10 | 2.01 | 0.30 |
| rs7177711 | 0.47 | 0.13 | 1.77 | 0.27 |
| rs936339 | 0.47 | 0.10 | 2.35 | 0.36 |
| rs994533 | 0.47 | 0.16 | 1.37 | 0.17 |
| rs429433 | 0.48 | 0.11 | 2.08 | 0.33 |
| rs11659752 | 0.49 | 0.14 | 1.71 | 0.26 |
| rs2377058 | 0.49 | 0.09 | 2.61 | 0.40 |
| rs17369123 | 0.50 | 0.16 | 1.57 | 0.24 |
| rs2735469 | 0.50 | 0.14 | 1.82 | 0.29 |
| rs6894139 | 0.50 | 0.20 | 1.25 | 0.14 |
| rs11616380 | 0.51 | 0.10 | 2.54 | 0.41 |
| rs26024 | 0.51 | 0.15 | 1.72 | 0.28 |
| rs11152213 | 0.53 | 0.15 | 1.94 | 0.34 |
| rs212524 | 0.53 | 0.14 | 1.98 | 0.34 |
| rs32855 | 0.53 | 0.14 | 2.07 | 0.36 |
| rs6974574 | 0.53 | 0.20 | 1.38 | 0.19 |
| rs1053996 | 0.54 | 0.13 | 2.24 | 0.40 |
| rs11640018 | 0.54 | 0.11 | 2.73 | 0.46 |
| rs12120956 | 0.54 | 0.15 | 1.98 | 0.35 |
| rs6921207 | 0.54 | 0.16 | 1.79 | 0.31 |
| rs782930 | 0.54 | 0.13 | 2.24 | 0.40 |
| rs8103068 | 0.55 | 0.15 | 1.98 | 0.36 |
| rs10997979 | 0.56 | 0.15 | 2.07 | 0.39 |
| rs1190545 | 0.57 | 0.17 | 1.93 | 0.36 |
| rs1599473 | 0.57 | 0.17 | 1.87 | 0.35 |
| rs3814333 | 0.57 | 0.31 | 1.04 | 0.07 |
| rs7853235 | 0.57 | 0.18 | 1.87 | 0.35 |
| rs1478610 | 0.58 | 0.16 | 2.05 | 0.40 |
| rs3885668 | 0.58 | 0.15 | 2.21 | 0.42 |
| rs7517682 | 0.58 | 0.17 | 1.90 | 0.36 |
| rs11100790 | 0.59 | 0.14 | 2.39 | 0.46 |
| rs12330322 | 0.59 | 0.22 | 1.58 | 0.29 |
| rs3116168 | 0.59 | 0.27 | 1.30 | 0.19 |
| rs4733724 | 0.59 | 0.31 | 1.16 | 0.13 |
| rs6540834 | 0.60 | 0.21 | 1.74 | 0.34 |
| rs8006657 | 0.60 | 0.15 | 2.35 | 0.46 |
| rs12669267 | 0.61 | 0.14 | 2.63 | 0.51 |
| rs7033487 | 0.61 | 0.25 | 1.52 | 0.29 |
| rs9650315 | 0.61 | 0.31 | 1.20 | 0.15 |
| rs2679184 | 0.62 | 0.21 | 1.89 | 0.40 |
| rs6085662 | 0.62 | 0.14 | 2.84 | 0.54 |
| rs7733195 | 0.63 | 0.23 | 1.67 | 0.35 |
| rs2013265 | 0.64 | 0.20 | 2.08 | 0.46 |
| rs2338115 | 0.64 | 0.19 | 2.14 | 0.47 |
| rs4785393 | 0.64 | 0.13 | 3.25 | 0.59 |
| rs1884897 | 0.65 | 0.34 | 1.24 | 0.19 |
| rs310421 | 0.65 | 0.27 | 1.52 | 0.32 |
| rs6485978 | 0.65 | 0.19 | 2.17 | 0.48 |
| rs1535466 | 0.66 | 0.21 | 2.06 | 0.48 |
| rs17038954 | 0.66 | 0.17 | 2.55 | 0.55 |
| rs181338 | 0.66 | 0.25 | 1.75 | 0.40 |
| rs2302580 | 0.66 | 0.25 | 1.70 | 0.39 |
| rs6061231 | 0.66 | 0.16 | 2.73 | 0.57 |
| rs6751657 | 0.66 | 0.21 | 2.03 | 0.47 |
| rs10152739 | 0.68 | 0.16 | 2.89 | 0.60 |
| rs13133465 | 0.68 | 0.15 | 3.01 | 0.61 |
| rs13416119 | 0.69 | 0.13 | 3.66 | 0.67 |
| rs4601530 | 0.69 | 0.20 | 2.45 | 0.57 |
| rs1659127 | 0.70 | 0.26 | 1.88 | 0.48 |
| rs1742829 | 0.70 | 0.18 | 2.67 | 0.60 |
| rs1950500 | 0.70 | 0.26 | 1.90 | 0.48 |
| rs3825199 | 0.70 | 0.37 | 1.34 | 0.28 |
| rs9858528 | 0.70 | 0.17 | 2.82 | 0.61 |
| rs2175513 | 0.71 | 0.14 | 3.59 | 0.68 |
| rs4246079 | 0.71 | 0.22 | 2.26 | 0.56 |
| rs2074977 | 0.72 | 0.27 | 1.93 | 0.51 |
| rs2326458 | 0.72 | 0.17 | 3.01 | 0.66 |
| rs11799609 | 0.73 | 0.18 | 2.95 | 0.66 |
| rs7654571 | 0.73 | 0.17 | 3.25 | 0.68 |
| rs16994718 | 0.74 | 0.15 | 3.74 | 0.71 |
| rs17410035 | 0.74 | 0.16 | 3.41 | 0.70 |
| rs17511102 | 0.74 | 0.26 | 2.13 | 0.58 |
| rs1171615 | 0.75 | 0.16 | 3.60 | 0.72 |
| rs12474201 | 0.75 | 0.27 | 2.05 | 0.57 |
| rs2829941 | 0.75 | 0.15 | 3.74 | 0.72 |
| rs7069985 | 0.75 | 0.18 | 3.09 | 0.69 |
| rs11624136 | 0.76 | 0.17 | 3.35 | 0.72 |
| rs1658351 | 0.76 | 0.22 | 2.65 | 0.67 |
| rs301901 | 0.76 | 0.25 | 2.34 | 0.63 |
| rs4656220 | 0.76 | 0.19 | 3.11 | 0.70 |
| rs6600365 | 0.76 | 0.28 | 2.05 | 0.58 |
| rs757081 | 0.76 | 0.23 | 2.54 | 0.66 |
| rs806794 | 0.77 | 0.47 | 1.25 | 0.29 |
| rs1199734 | 0.78 | 0.15 | 4.07 | 0.77 |
| rs3118905 | 0.78 | 0.47 | 1.30 | 0.34 |
| rs3828760 | 0.78 | 0.19 | 3.21 | 0.73 |
| rs2079795 | 0.79 | 0.41 | 1.53 | 0.48 |
| rs2390151 | 0.79 | 0.27 | 2.35 | 0.67 |
| rs11722554 | 0.80 | 0.22 | 2.91 | 0.74 |
| rs17450430 | 0.80 | 0.31 | 2.05 | 0.64 |
| rs7154721 | 0.80 | 0.30 | 2.17 | 0.67 |
| rs7568069 | 0.80 | 0.23 | 2.76 | 0.72 |
| rs8180991 | 0.80 | 0.24 | 2.60 | 0.70 |
| rs9841435 | 0.80 | 0.19 | 3.41 | 0.76 |
| rs11049611 | 0.81 | 0.37 | 1.77 | 0.60 |
| rs12519505 | 0.81 | 0.16 | 4.01 | 0.79 |
| rs13388725 | 0.81 | 0.17 | 3.93 | 0.79 |
| rs3760318 | 0.82 | 0.41 | 1.61 | 0.56 |
| rs6457374 | 0.82 | 0.39 | 1.73 | 0.61 |
| rs7162825 | 0.83 | 0.16 | 4.41 | 0.83 |
| rs11244750 | 0.84 | 0.16 | 4.29 | 0.83 |
| rs6544089 | 0.84 | 0.20 | 3.46 | 0.80 |
| rs12119525 | 0.85 | 0.17 | 4.31 | 0.85 |
| rs6838153 | 0.85 | 0.23 | 3.10 | 0.80 |
| rs7659107 | 0.85 | 0.20 | 3.56 | 0.82 |
| rs4896582 | 0.86 | 0.48 | 1.55 | 0.62 |
| rs10948222 | 0.88 | 0.36 | 2.14 | 0.77 |
| rs2811594 | 0.89 | 0.28 | 2.85 | 0.85 |
| rs1036821 | 0.90 | 0.41 | 1.99 | 0.79 |
| rs10766065 | 0.90 | 0.18 | 4.43 | 0.90 |
| rs12323101 | 0.90 | 0.23 | 3.49 | 0.88 |
| rs2737220 | 0.90 | 0.18 | 4.52 | 0.90 |
| rs6794009 | 0.90 | 0.16 | 5.00 | 0.90 |
| rs7043114 | 0.90 | 0.34 | 2.37 | 0.82 |
| rs33852 | 0.91 | 0.34 | 2.43 | 0.85 |
| rs7716219 | 0.91 | 0.34 | 2.42 | 0.85 |
| rs12137162 | 0.92 | 0.18 | 4.72 | 0.92 |
| rs1325596 | 0.92 | 0.30 | 2.78 | 0.88 |
| rs8017130 | 0.92 | 0.25 | 3.41 | 0.90 |
| rs1935157 | 0.93 | 0.27 | 3.12 | 0.90 |
| rs34651 | 0.93 | 0.26 | 3.32 | 0.91 |
| rs3791679 | 0.93 | 0.54 | 1.58 | 0.78 |
| rs9993613 | 0.93 | 0.37 | 2.33 | 0.87 |
| rs12882130 | 0.94 | 0.29 | 3.06 | 0.92 |
| rs12411277 | 0.95 | 0.25 | 3.57 | 0.94 |
| rs798497 | 0.96 | 0.57 | 1.61 | 0.87 |
| rs16964211 | 0.97 | 0.34 | 2.79 | 0.96 |
| rs2211866 | 0.97 | 0.28 | 3.42 | 0.97 |
| rs7334755 | 0.98 | 0.33 | 2.90 | 0.97 |
| rs9434723 | 0.98 | 0.26 | 3.77 | 0.98 |
| rs17349981 | 0.99 | 0.19 | 5.17 | 0.99 |
| rs3812591 | 0.99 | 0.28 | 3.45 | 0.98 |
| rs7849585 | 0.99 | 0.44 | 2.23 | 0.97 |
| rs11633371 | 1.00 | 0.38 | 2.66 | 1.00 |
| rs12538407 | 1.00 | 0.42 | 2.38 | 0.99 |
| rs13393800 | 1.00 | 0.31 | 3.23 | 1.00 |
| rs2581830 | 1.00 | 0.41 | 2.40 | 0.99 |
| rs3802758 | 1.00 | 0.26 | 3.78 | 1.00 |
| rs6563199 | 1.00 | 0.21 | 4.85 | 1.00 |
| rs9428104 | 1.00 | 0.48 | 2.07 | 0.99 |
| rs2028067 | 1.01 | 0.30 | 3.36 | 0.99 |
| rs10059761 | 1.02 | 0.20 | 5.16 | 0.99 |
| rs12615742 | 1.02 | 0.30 | 3.47 | 0.97 |
| rs7823327 | 1.02 | 0.23 | 4.53 | 0.98 |
| rs891088 | 1.02 | 0.35 | 2.99 | 0.97 |
| rs10962832 | 1.03 | 0.21 | 5.06 | 0.97 |
| rs17330192 | 1.03 | 0.20 | 5.27 | 0.97 |
| rs4246302 | 1.03 | 0.34 | 3.09 | 0.96 |
| rs1681630 | 1.04 | 0.37 | 2.91 | 0.94 |
| rs2164968 | 1.05 | 0.22 | 5.10 | 0.95 |
| rs2531992 | 1.05 | 0.26 | 4.28 | 0.94 |
| rs1113765 | 1.06 | 0.23 | 4.90 | 0.94 |
| rs1346490 | 1.06 | 0.22 | 5.21 | 0.94 |
| rs4425077 | 1.06 | 0.26 | 4.27 | 0.93 |
| rs833706 | 1.07 | 0.22 | 5.27 | 0.93 |
| rs4256170 | 1.08 | 0.66 | 1.78 | 0.76 |
| rs6696239 | 1.08 | 0.44 | 2.67 | 0.86 |
| rs17081935 | 1.09 | 0.37 | 3.26 | 0.88 |
| rs6441170 | 1.09 | 0.30 | 3.90 | 0.89 |
| rs13078528 | 1.10 | 0.28 | 4.40 | 0.89 |
| rs17556750 | 1.11 | 0.58 | 2.16 | 0.75 |
| rs3958122 | 1.11 | 0.38 | 3.26 | 0.84 |
| rs1546391 | 1.12 | 0.29 | 4.25 | 0.87 |
| rs2306694 | 1.12 | 0.34 | 3.67 | 0.85 |
| rs1529701 | 1.13 | 0.28 | 4.55 | 0.86 |
| rs13006748 | 1.18 | 0.32 | 4.44 | 0.80 |
| rs7692995 | 1.18 | 0.71 | 1.98 | 0.52 |
| rs1996422 | 1.19 | 0.30 | 4.69 | 0.81 |
| rs10794175 | 1.20 | 0.31 | 4.74 | 0.79 |
| rs12214804 | 1.20 | 0.67 | 2.15 | 0.54 |
| rs1815314 | 1.20 | 0.34 | 4.31 | 0.77 |
| rs4686904 | 1.20 | 0.31 | 4.64 | 0.79 |
| rs552707 | 1.20 | 0.63 | 2.25 | 0.58 |
| rs6879260 | 1.20 | 0.41 | 3.52 | 0.74 |
| rs2034172 | 1.21 | 0.23 | 6.25 | 0.82 |
| rs2120335 | 1.21 | 0.27 | 5.34 | 0.80 |
| rs3763631 | 1.21 | 0.26 | 5.71 | 0.81 |
| rs4350272 | 1.21 | 0.25 | 5.74 | 0.81 |
| rs16968242 | 1.22 | 0.25 | 5.99 | 0.81 |
| rs606452 | 1.22 | 0.48 | 3.08 | 0.68 |
| rs1155939 | 1.23 | 0.65 | 2.34 | 0.53 |
| rs17792664 | 1.24 | 0.33 | 4.63 | 0.75 |
| rs9395264 | 1.24 | 0.29 | 5.34 | 0.77 |
| rs955748 | 1.25 | 0.40 | 3.85 | 0.70 |
| rs17603945 | 1.26 | 0.41 | 3.86 | 0.68 |
| rs291979 | 1.26 | 0.41 | 3.87 | 0.69 |
| rs6691924 | 1.26 | 0.30 | 5.40 | 0.75 |
| rs8102380 | 1.27 | 0.29 | 5.58 | 0.75 |
| rs2072268 | 1.28 | 0.32 | 5.21 | 0.73 |
| rs6439168 | 1.29 | 0.49 | 3.38 | 0.61 |
| rs6920372 | 1.30 | 0.42 | 4.00 | 0.64 |
| rs9816693 | 1.30 | 0.40 | 4.22 | 0.66 |
| rs2289195 | 1.31 | 0.62 | 2.81 | 0.48 |
| rs9217 | 1.31 | 0.48 | 3.58 | 0.60 |
| rs4802134 | 1.32 | 0.42 | 4.19 | 0.64 |
| rs11880992 | 1.33 | 0.57 | 3.10 | 0.50 |
| rs7701414 | 1.33 | 0.64 | 2.77 | 0.44 |
| rs3915129 | 1.36 | 0.24 | 7.64 | 0.73 |
| rs6658763 | 1.36 | 0.33 | 5.71 | 0.67 |
| rs915506 | 1.36 | 0.33 | 5.58 | 0.67 |
| rs6137287 | 1.37 | 0.30 | 6.32 | 0.69 |
| rs2573625 | 1.38 | 0.49 | 3.85 | 0.54 |
| rs2166898 | 1.39 | 0.32 | 6.03 | 0.66 |
| rs11648796 | 1.40 | 0.50 | 3.91 | 0.52 |
| rs2854207 | 1.40 | 0.73 | 2.68 | 0.31 |
| rs425277 | 1.40 | 0.47 | 4.20 | 0.55 |
| rs9392918 | 1.41 | 0.69 | 2.88 | 0.35 |
| rs1074683 | 1.42 | 0.69 | 2.93 | 0.34 |
| rs2280470 | 1.42 | 0.70 | 2.85 | 0.33 |
| rs6584575 | 1.42 | 0.37 | 5.41 | 0.61 |
| rs10283100 | 1.43 | 0.44 | 4.64 | 0.55 |
| rs4953951 | 1.43 | 0.38 | 5.48 | 0.60 |
| rs724016 | 1.43 | 1.02 | 2.02 | 0.04 |
| rs991967 | 1.43 | 0.59 | 3.46 | 0.43 |
| rs1244981 | 1.44 | 0.29 | 7.32 | 0.66 |
| rs3739707 | 1.44 | 0.37 | 5.55 | 0.59 |
| rs7712162 | 1.44 | 0.32 | 6.51 | 0.64 |
| rs17777628 | 1.45 | 0.37 | 5.77 | 0.60 |
| rs2631676 | 1.45 | 0.42 | 5.06 | 0.56 |
| rs8042424 | 1.45 | 0.36 | 5.85 | 0.60 |
| rs9880211 | 1.45 | 0.51 | 4.09 | 0.49 |
| rs568610 | 1.46 | 0.35 | 6.20 | 0.60 |
| rs11750568 | 1.47 | 0.33 | 6.45 | 0.61 |
| rs11618507 | 1.48 | 0.37 | 5.98 | 0.58 |
| rs4735677 | 1.48 | 0.66 | 3.35 | 0.34 |
| rs11687941 | 1.50 | 0.42 | 5.36 | 0.53 |
| rs2345835 | 1.50 | 0.32 | 7.12 | 0.61 |
| rs7646824 | 1.50 | 0.28 | 8.20 | 0.64 |
| rs1562975 | 1.52 | 0.44 | 5.19 | 0.51 |
| rs2781373 | 1.52 | 0.39 | 6.00 | 0.55 |
| rs7466269 | 1.52 | 0.63 | 3.67 | 0.35 |
| rs227724 | 1.54 | 0.55 | 4.36 | 0.41 |
| rs4725061 | 1.55 | 0.38 | 6.31 | 0.54 |
| rs4973429 | 1.55 | 0.59 | 4.11 | 0.37 |
| rs3020418 | 1.56 | 0.62 | 3.97 | 0.35 |
| rs6955948 | 1.56 | 0.61 | 4.00 | 0.35 |
| rs4369779 | 1.57 | 0.85 | 2.87 | 0.15 |
| rs4620037 | 1.57 | 0.53 | 4.67 | 0.42 |
| rs12533079 | 1.58 | 0.35 | 7.07 | 0.55 |
| rs2597513 | 1.58 | 0.48 | 5.19 | 0.45 |
| rs992157 | 1.58 | 0.45 | 5.56 | 0.47 |
| rs1321666 | 1.59 | 0.32 | 7.90 | 0.57 |
| rs314263 | 1.59 | 0.82 | 3.08 | 0.17 |
| rs4986172 | 1.60 | 0.69 | 3.69 | 0.27 |
| rs7253628 | 1.61 | 0.35 | 7.38 | 0.54 |
| rs16834765 | 1.63 | 0.41 | 6.39 | 0.49 |
| rs4320932 | 1.65 | 0.51 | 5.38 | 0.41 |
| rs7870753 | 1.65 | 0.78 | 3.49 | 0.19 |
| rs7126398 | 1.66 | 0.45 | 6.07 | 0.44 |
| rs9327705 | 1.66 | 0.33 | 8.35 | 0.54 |
| rs1326023 | 1.67 | 0.49 | 5.64 | 0.41 |
| rs14062 | 1.68 | 0.33 | 8.40 | 0.53 |
| rs6761041 | 1.68 | 0.52 | 5.51 | 0.39 |
| rs8058684 | 1.68 | 0.41 | 6.88 | 0.47 |
| rs2093210 | 1.70 | 0.83 | 3.44 | 0.14 |
| rs42039 | 1.70 | 1.07 | 2.69 | 0.02 |
| rs738288 | 1.70 | 0.42 | 6.83 | 0.46 |
| rs7561273 | 1.70 | 0.57 | 5.00 | 0.34 |
| rs39623 | 1.72 | 0.39 | 7.65 | 0.48 |
| rs2815379 | 1.73 | 0.32 | 9.37 | 0.52 |
| rs564914 | 1.73 | 0.53 | 5.59 | 0.36 |
| rs354196 | 1.74 | 0.47 | 6.42 | 0.41 |
| rs932445 | 1.74 | 0.36 | 8.54 | 0.49 |
| rs3812423 | 1.78 | 0.46 | 6.89 | 0.40 |
| rs1797625 | 1.79 | 0.41 | 7.84 | 0.44 |
| rs11855014 | 1.81 | 0.47 | 6.96 | 0.39 |
| rs584828 | 1.81 | 0.57 | 5.75 | 0.31 |
| rs1966913 | 1.82 | 0.38 | 8.66 | 0.45 |
| rs6919534 | 1.82 | 0.81 | 4.06 | 0.14 |
| rs4332428 | 1.88 | 0.58 | 6.09 | 0.29 |
| rs1233627 | 1.90 | 0.63 | 5.77 | 0.26 |
| rs3014219 | 1.91 | 0.53 | 6.86 | 0.32 |
| rs10767838 | 1.93 | 0.56 | 6.60 | 0.30 |
| rs12914466 | 1.94 | 0.35 | 10.83 | 0.45 |
| rs6694089 | 1.95 | 0.89 | 4.24 | 0.09 |
| rs862034 | 1.97 | 0.71 | 5.48 | 0.19 |
| rs12470505 | 1.98 | 0.74 | 5.27 | 0.17 |
| rs2023693 | 1.98 | 0.37 | 10.53 | 0.42 |
| rs4834927 | 1.98 | 0.41 | 9.61 | 0.40 |
| rs4843367 | 1.98 | 0.42 | 9.41 | 0.39 |
| rs10883563 | 1.99 | 0.60 | 6.61 | 0.26 |
| rs2298265 | 1.99 | 0.45 | 8.70 | 0.36 |
| rs10152591 | 2.00 | 0.68 | 5.89 | 0.21 |
| rs11661645 | 2.02 | 0.45 | 9.15 | 0.36 |
| rs4624820 | 2.02 | 0.46 | 8.90 | 0.35 |
| rs9409082 | 2.02 | 0.68 | 5.98 | 0.20 |
| rs1832871 | 2.03 | 0.64 | 6.48 | 0.23 |
| rs3782089 | 2.04 | 0.73 | 5.73 | 0.18 |
| rs6446315 | 2.04 | 0.53 | 7.94 | 0.30 |
| rs6949739 | 2.05 | 0.52 | 8.03 | 0.30 |
| rs11634405 | 2.06 | 0.49 | 8.72 | 0.32 |
| rs8103992 | 2.08 | 0.62 | 6.99 | 0.23 |
| rs8097893 | 2.10 | 0.41 | 10.81 | 0.37 |
| rs10995319 | 2.11 | 0.38 | 11.70 | 0.39 |
| rs1401795 | 2.11 | 0.79 | 5.66 | 0.14 |
| rs2856321 | 2.12 | 0.86 | 5.24 | 0.10 |
| rs9766 | 2.12 | 0.60 | 7.55 | 0.25 |
| rs165189 | 2.13 | 0.53 | 8.62 | 0.29 |
| rs9825951 | 2.13 | 0.58 | 7.81 | 0.26 |
| rs8052560 | 2.20 | 0.84 | 5.74 | 0.11 |
| rs9291926 | 2.21 | 0.53 | 9.19 | 0.27 |
| rs2974438 | 2.22 | 0.87 | 5.67 | 0.10 |
| rs5757318 | 2.22 | 0.58 | 8.51 | 0.25 |
| rs11684404 | 2.25 | 0.94 | 5.39 | 0.07 |
| rs13150868 | 2.33 | 0.45 | 12.13 | 0.31 |
| rs9977276 | 2.34 | 0.53 | 10.36 | 0.26 |
| rs1047014 | 2.37 | 0.85 | 6.63 | 0.10 |
| rs12855 | 2.38 | 0.68 | 8.30 | 0.17 |
| rs6020202 | 2.38 | 0.53 | 10.64 | 0.26 |
| rs867245 | 2.39 | 0.56 | 10.27 | 0.24 |
| rs6080830 | 2.44 | 0.41 | 14.44 | 0.32 |
| rs2378870 | 2.47 | 0.58 | 10.54 | 0.22 |
| rs8117259 | 2.47 | 0.49 | 12.58 | 0.28 |
| rs991946 | 2.48 | 0.65 | 9.52 | 0.18 |
| rs2806561 | 2.50 | 0.88 | 7.10 | 0.09 |
| rs3800461 | 2.54 | 1.20 | 5.39 | 0.01 |
| rs6952113 | 2.54 | 0.55 | 11.81 | 0.23 |
| rs12513181 | 2.59 | 0.55 | 12.16 | 0.23 |
| rs17250196 | 2.62 | 0.68 | 10.10 | 0.16 |
| rs11616067 | 2.63 | 0.53 | 12.97 | 0.24 |
| rs509035 | 2.63 | 1.05 | 6.62 | 0.04 |
| rs12693589 | 2.66 | 0.63 | 11.25 | 0.18 |
| rs4273857 | 2.67 | 0.82 | 8.65 | 0.10 |
| rs11236294 | 2.68 | 0.52 | 13.80 | 0.24 |
| rs12209223 | 2.71 | 1.12 | 6.55 | 0.03 |
| rs692964 | 2.72 | 0.60 | 12.26 | 0.19 |
| rs1405212 | 2.73 | 0.81 | 9.23 | 0.11 |
| rs3129254 | 2.76 | 0.62 | 12.42 | 0.18 |
| rs13177718 | 2.82 | 0.84 | 9.53 | 0.09 |
| rs7273787 | 2.88 | 0.74 | 11.17 | 0.13 |
| rs958225 | 2.88 | 0.72 | 11.46 | 0.13 |
| rs389663 | 2.91 | 0.75 | 11.27 | 0.12 |
| rs2509133 | 2.94 | 0.62 | 13.97 | 0.17 |
| rs2633761 | 2.97 | 0.55 | 16.09 | 0.21 |
| rs318095 | 3.02 | 0.98 | 9.31 | 0.05 |
| rs989393 | 3.07 | 0.75 | 12.56 | 0.12 |
| rs12186664 | 3.08 | 0.74 | 12.71 | 0.12 |
| rs2834442 | 3.13 | 0.97 | 10.15 | 0.06 |
| rs4812586 | 3.17 | 0.93 | 10.82 | 0.07 |
| rs1625895 | 3.18 | 0.81 | 12.54 | 0.10 |
| rs711245 | 3.21 | 0.94 | 10.93 | 0.06 |
| rs8067165 | 3.22 | 0.94 | 11.08 | 0.06 |
| rs209918 | 3.33 | 0.76 | 14.56 | 0.11 |
| rs2275325 | 3.34 | 0.67 | 16.69 | 0.14 |
| rs2058092 | 3.36 | 0.65 | 17.47 | 0.15 |
| rs1014987 | 3.47 | 0.71 | 16.99 | 0.12 |
| rs3812040 | 3.49 | 0.95 | 12.79 | 0.06 |
| rs7284476 | 3.51 | 0.72 | 17.02 | 0.12 |
| rs11867943 | 3.64 | 0.77 | 17.22 | 0.10 |
| rs2057291 | 3.69 | 0.81 | 16.84 | 0.09 |
| rs7319045 | 3.96 | 1.25 | 12.51 | 0.02 |
| rs8073371 | 3.99 | 1.09 | 14.55 | 0.04 |
| rs2219320 | 4.11 | 0.99 | 17.10 | 0.05 |
| rs7567851 | 4.12 | 1.17 | 14.47 | 0.03 |
| rs6988484 | 4.15 | 1.02 | 16.80 | 0.05 |
| rs10083886 | 4.19 | 0.86 | 20.29 | 0.08 |
| rs17807185 | 4.19 | 1.12 | 15.65 | 0.03 |
| rs17659078 | 4.48 | 1.06 | 18.87 | 0.04 |
| rs10119624 | 4.60 | 1.41 | 15.02 | 0.01 |
| rs17163588 | 4.83 | 1.30 | 17.96 | 0.02 |
| rs12863103 | 4.90 | 1.01 | 23.75 | 0.05 |
| rs13113518 | 5.95 | 1.25 | 28.23 | 0.02 |
| rs17806888 | 6.45 | 1.78 | 23.34 | 0.005 |
| rs12190423 | 6.53 | 1.26 | 33.96 | 0.03 |
| rs11880124 | 7.78 | 2.38 | 25.45 | 6.96×10^-4^ |
| rs6829680 | 8.66 | 1.74 | 43.01 | 0.01 |
| rs2306596 | 9.22 | 2.20 | 38.67 | 0.002 |
| rs4357716 | 12.46 | 3.52 | 44.13 | 9.23×10^-5^ |
| rs11156098 | 12.78 | 2.65 | 61.77 | 0.002 |

**Table S13. The outlying SNPs identified by MR-PRESSO for the risk factors of overall prostate cancer**

| **Risk factor** | **SNP** | **CHR** | **Position** | **Gene** |
| --- | --- | --- | --- | --- |
| Alcohol | rs13032049 | 2 | 63581507 | WDPCP |
| Alcohol | rs13094887 | 3 | 70968431 | Unknown |
| Alcohol | rs1713676 | 11 | 113660576 | Unknown |
| Alcohol | rs2011092 | 3 | 141124607 | ZBTB38 |
| Alcohol | rs28929474 | 14 | 94844947 | SERPINA1 |
| Alcohol | rs56030824 | 11 | 47397353 | SPI1 |
| Alcohol | rs823114 | 1 | 205719532 | Unknown |
| BMI | rs10971712 | 9 | 33820940 | UBE2R2 |
| BMI | rs11629783 | 15 | 66449049 | MAP2K1 |
| BMI | rs17636031 | 10 | 124905509 | Unknown |
| BMI | rs2185027 | 6 | 153060487 | RGS17 |
| BMI | rs2814992 | 6 | 34649367 | C6orf106 |
| BMI | rs905938 | 1 | 155018913 | ZBTB7B |
| Birth weight | rs72851023 | 11 | 2109390 | IGF2/H19 |
| Birth weight | rs7402982 | 15 | 98650040 | *IGF1R* |
| Birth weight | rs900399 | 3 | 157080943 | LINC02029 |
| Height | rs11880124 | 19 | 47375631 | DEDD2 |
| Height | rs12639764 | 4 | 106435654 | TET2 |
| Height | rs1625895 | 17 | 7518840 | TP53 |
| Height | rs17659078 | 11 | 2241166 | ASCL2 |
| Height | rs17783015 | 12 | 88755517 | ATP2B1 |
| Height | rs3118905 | 13 | 50003335 | DLEU7 |
| Height | rs3807931 | 7 | 20348199 | ITGB8 |
| Height | rs4344931 | 2 | 241467200 | AGXT |
| Height | rs4357716 | 11 | 68872342 | MYEOV |
| Height | rs4803468 | 19 | 46614192 | BCKDHA |
| Height | rs724016 | 3 | 142588260 | ZBTB38 |
| Height | rs7899004 | 10 | 104331425 | SUFU |
| Height | rs9309101 | 2 | 43483116 | THADA |
| Height | rs960006 | 16 | 4851196 | UBN1 |
| Sugar/Sucrose (glucose) | rs11715915 | 3 | 49417897 | AMT |
| Total fat | rs1260326 | 2 | 27508073 | GCKR |
| Total fat | rs429358 | 19 | 44908684 | APOE |
| Waist circumference | rs16894959 | 6 | 34933640 | UHRF1BP1 |
| Waist circumference | rs2075650 | 19 | 50087459 | TOMM40 |
| Waist circumference | rs2293576 | 11 | 47391562 | SLC39A13 |
| Waist circumference | rs6163 | 10 | 104586914 | CYP17A1 |
| Waist circumference | rs6440003 | 3 | 142576899 | ZBTB38 |
| WHR | rs2075650 | 19 | 50087459 | TOMM40 |

**Table S14. The outlying SNPs identified by MR-PRESSO for the risk factors of aggressive prostate cancer.**

| **Risk factor** | **SNP** | **CHR** | **Position** | **Gene** |
| --- | --- | --- | --- | --- |
| Alcohol | rs13032049 | 2 | 63581507 | WDPCP |
| Alcohol | rs823114 | 1 | 205719532 | Unknown |
| BMI | rs2185027 | 6 | 153060487 | RGS17 |
| BMI | rs2814992 | 6 | 34649367 | C6orf106 |
| BMI | rs7556169 | 1 | 8681342 | RERE |
| Birth weight | rs72851023 | 11 | 2109390 | IGF2/H19 |
| Height | rs7899004 | 10 | 104,331,425 | SUFU |
| Waist circumference | rs16894959 | 6 | 34933640 | UHRF1BP1 |

**Figure S1. Results of Mendelian randomization sensitivity analyses performed for overall prostate cancer risk.**

| 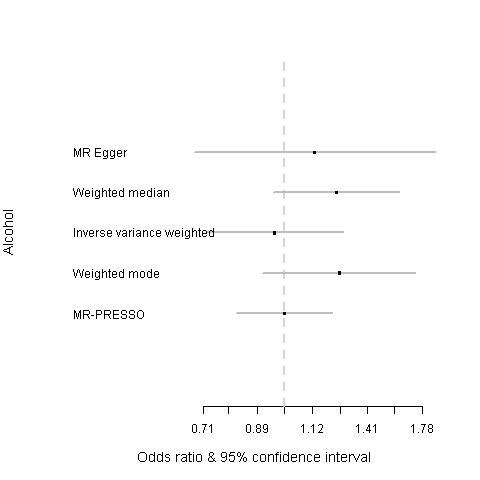 | 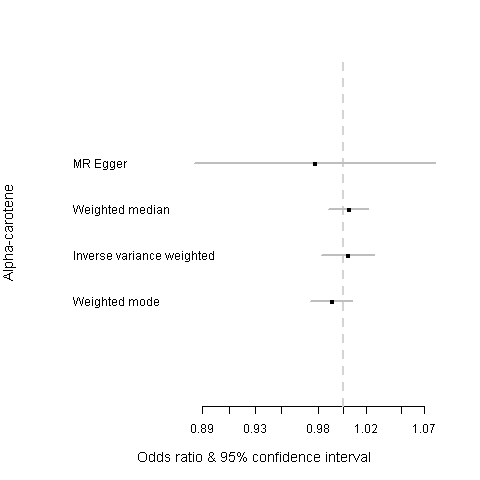 |
| --- | --- |
| 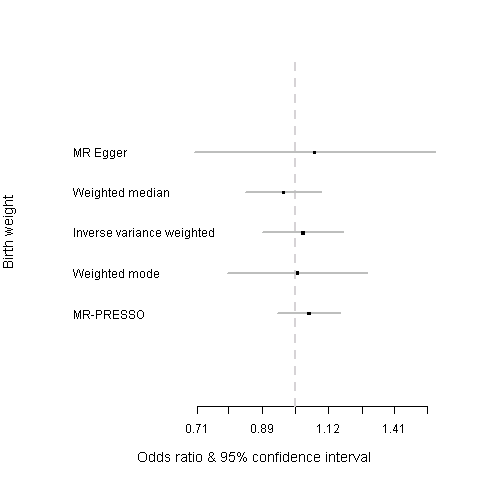 | 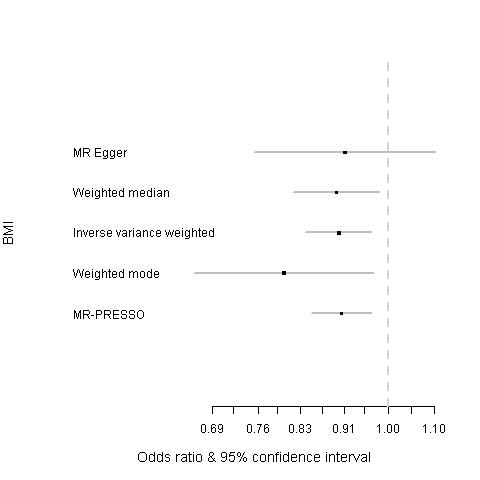 |
| 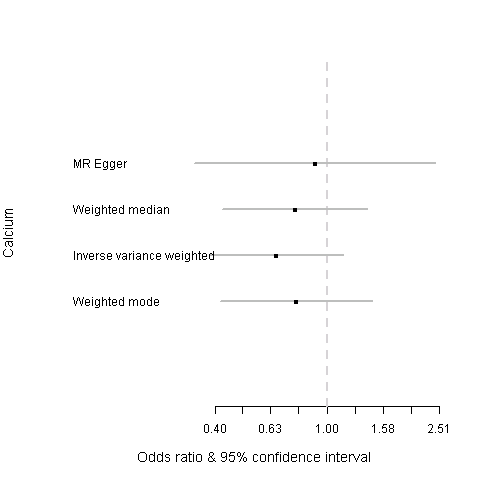 | 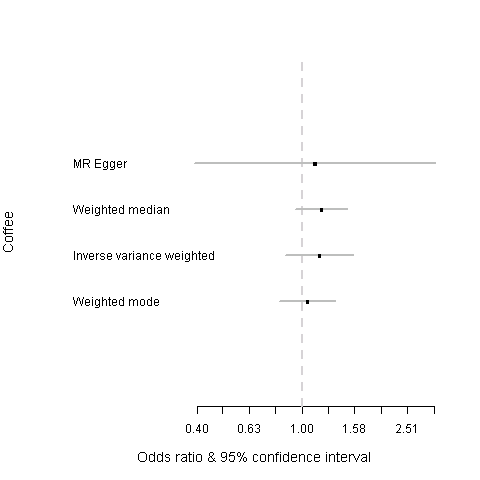 |
| 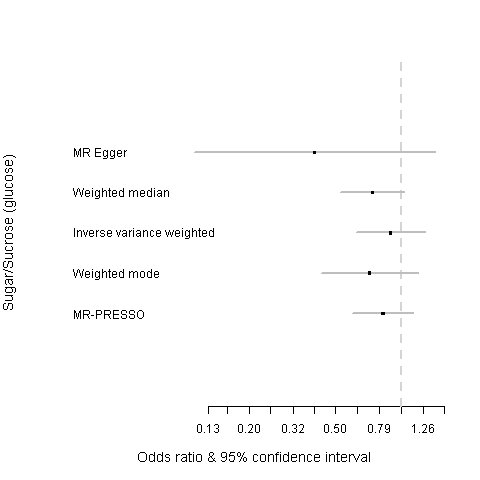 | 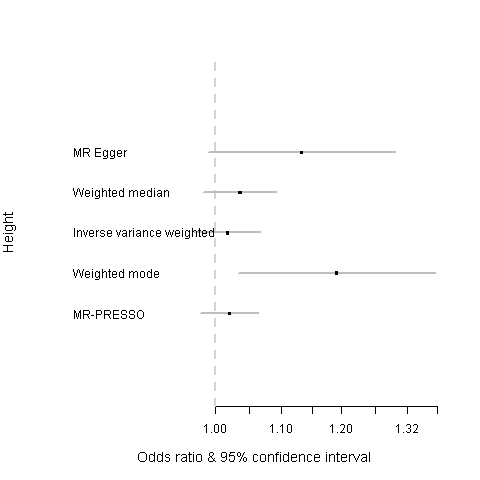 |
| 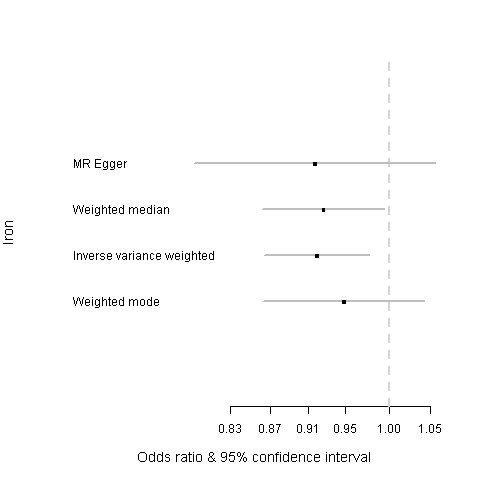 | 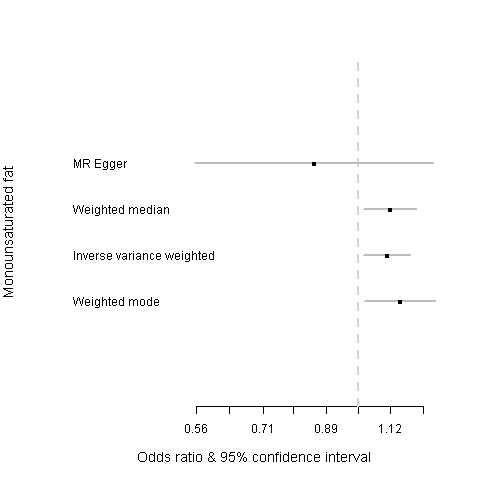 |
| 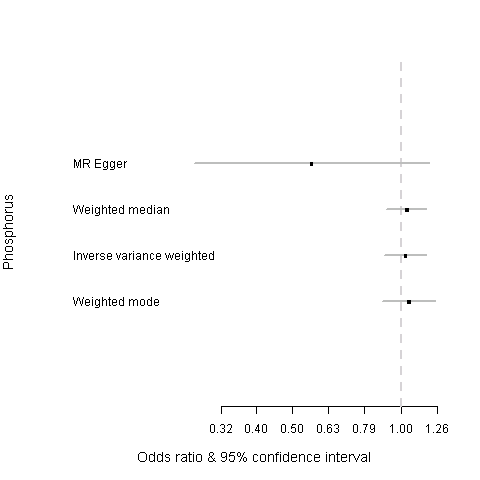 | 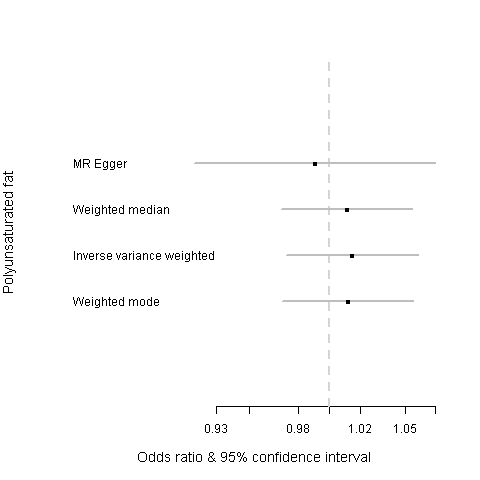 |
| 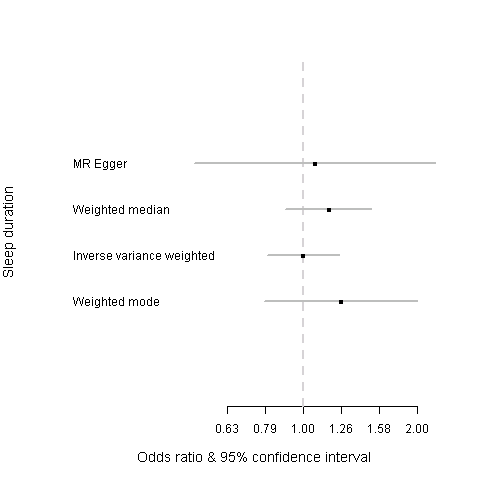 | 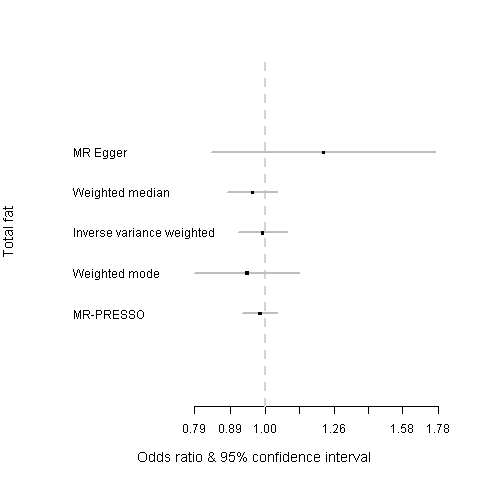 |
| 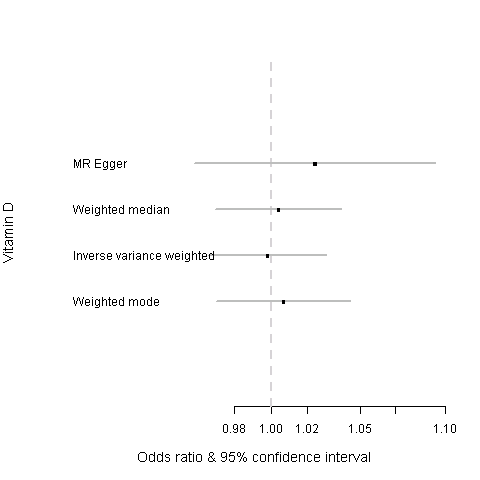 | 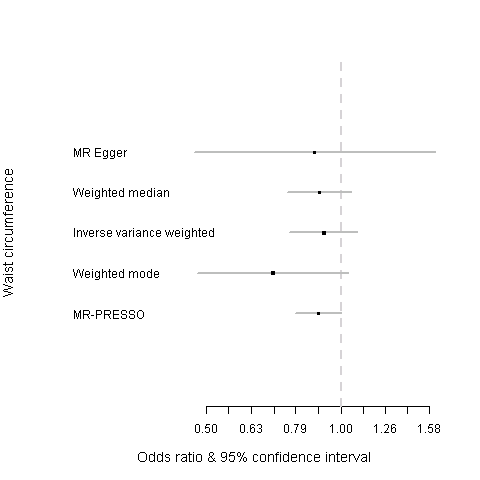 |
| 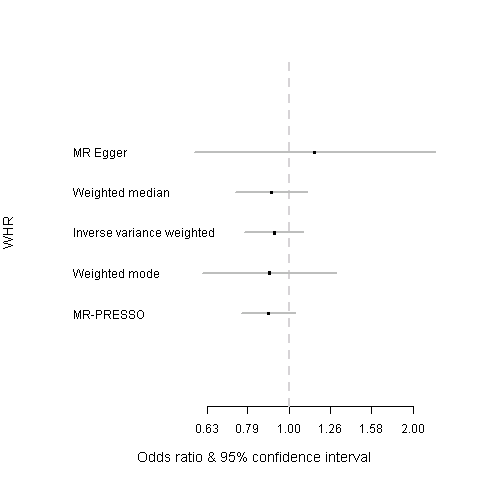 |  |

MR-PRESSO = when MR-PRESSO identified outlying SNPs for a risk factor, IVW method was re-ran after excluding these outlying SNPs.

**Figure S2. Results from MR-Base showing association between iron and overall prostate cancer risk A) comparison of results using different MR methods; B) funnel plot of IVW and MR-Egger regression**

| 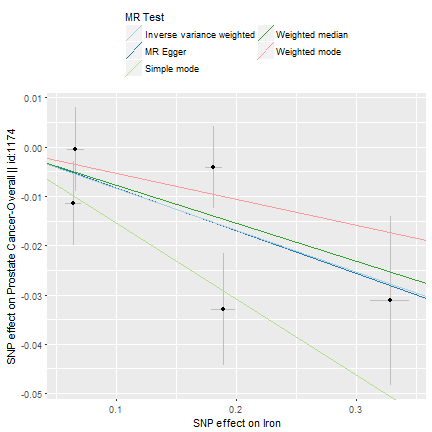**A)** | 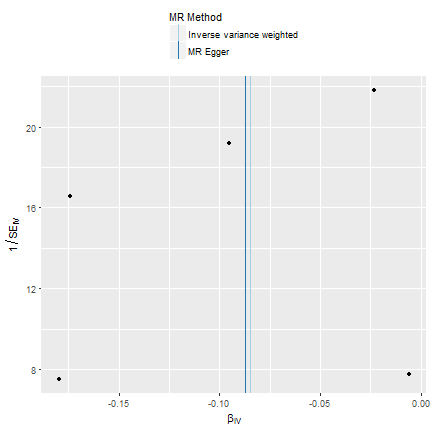**B)** |
| --- | --- |

**Figure S3. Results from MR-Base showing association between BMI and overall prostate cancer risk A) comparison of results using different MR methods; B) funnel plot of IVW and MR-Egger regression**

| 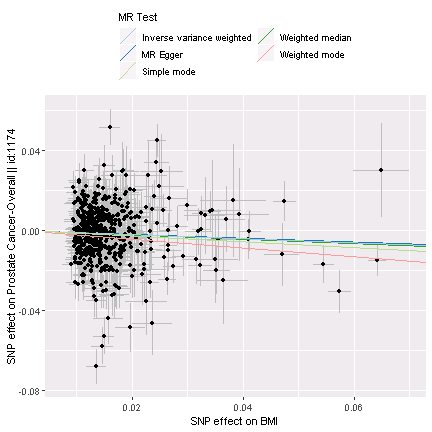**A)** | 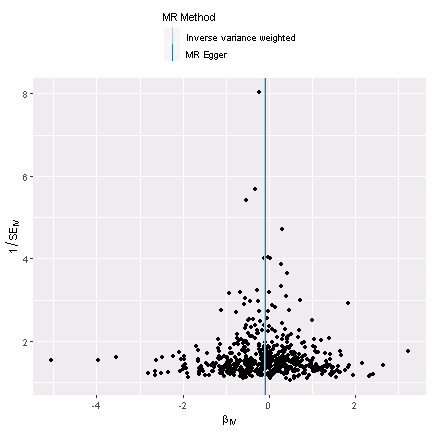**B)** |
| --- | --- |

**Figure S4. Results from MR-Base showing association between mono-unsaturated fat and overall prostate cancer risk A) comparison of results using different MR methods; B) funnel plot of IVW and MR-Egger regression**

| 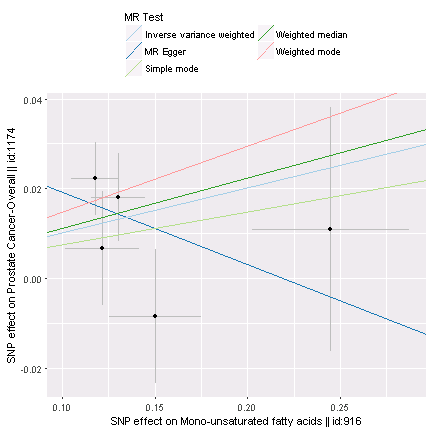**A)** | 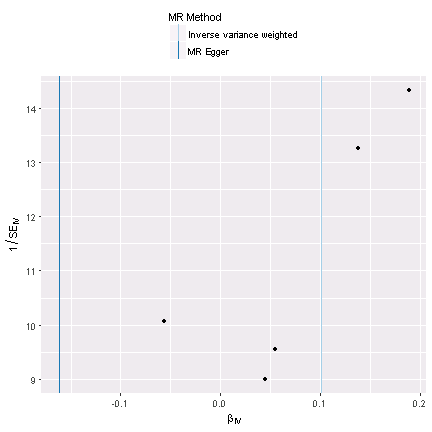**B)** |
| --- | --- |

**Figure S5. Results from MR-Base showing association between height and aggressive prostate cancer risk A) comparison of results using different MR methods; B) funnel plot of IVW and MR-Egger regression**

| 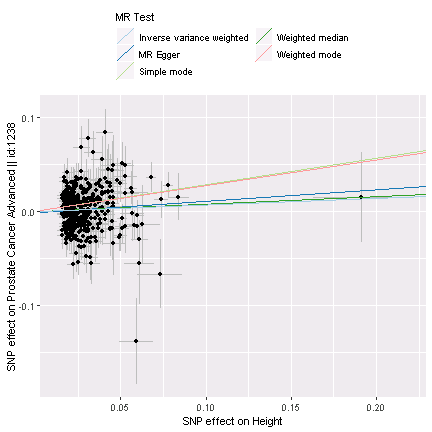**A)** | 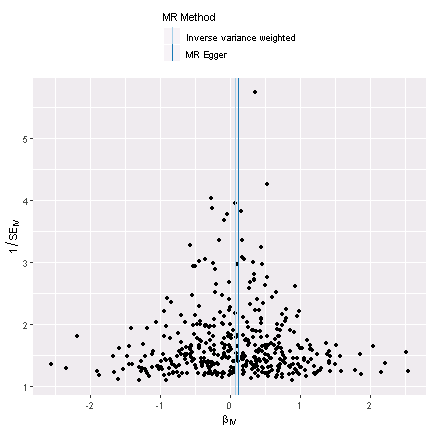**B)** |
| --- | --- |

**Figure S6. Results of Mendelian randomization sensitivity analyses performed for aggressive prostate cancer risk.**

| 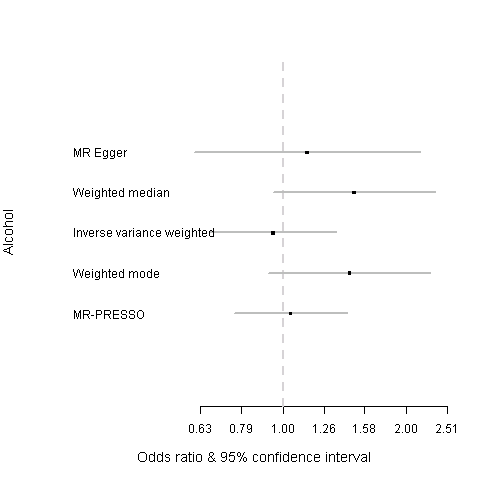 | 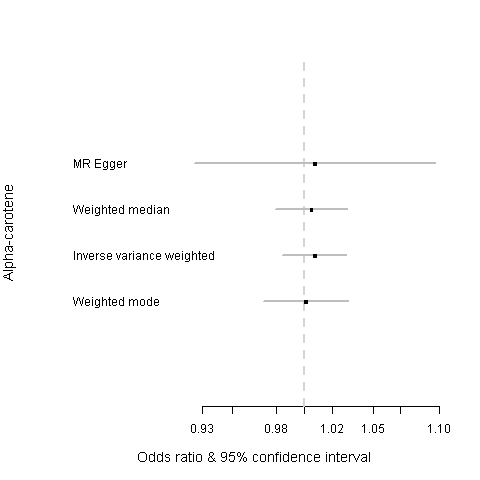 |
| --- | --- |
| 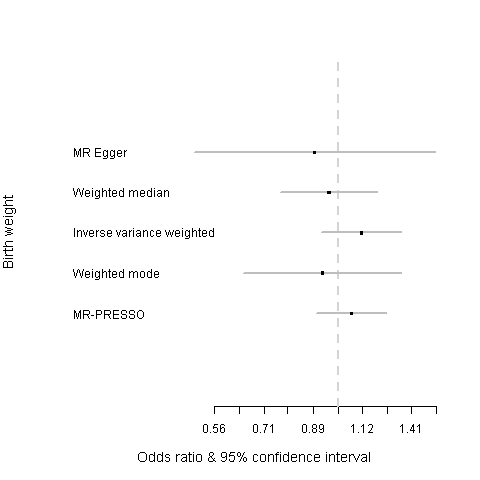 | 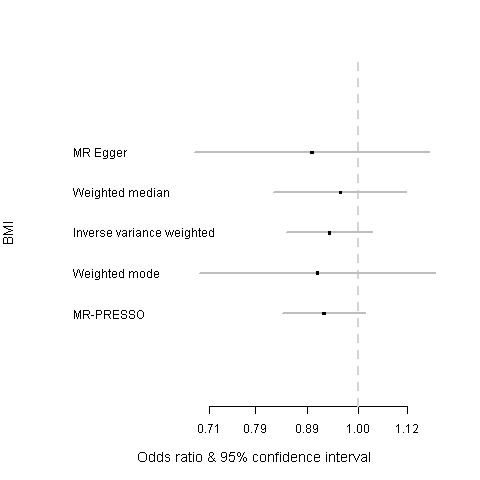 |
| 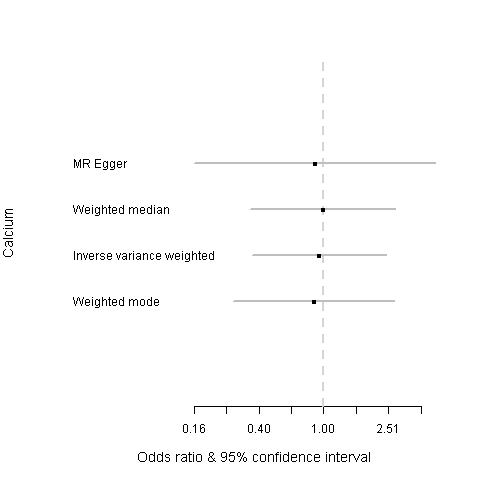 | 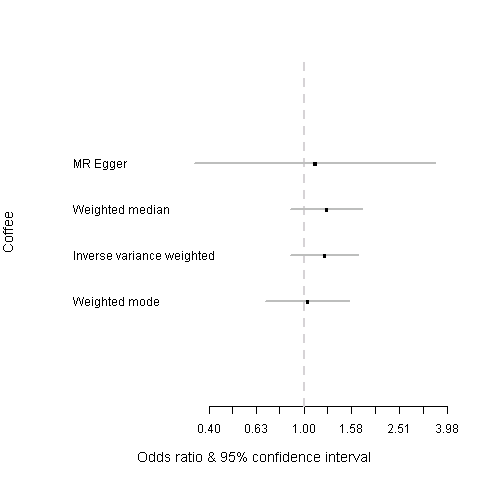 |
| 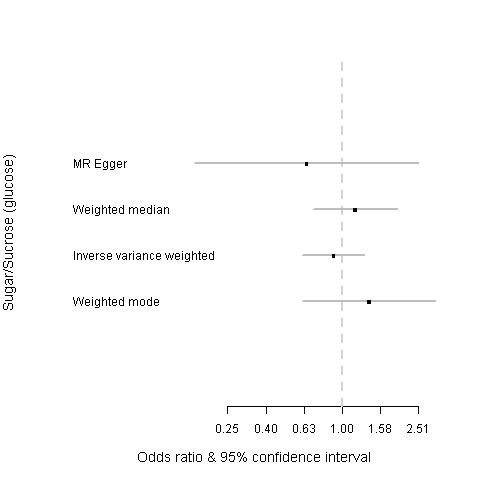 | 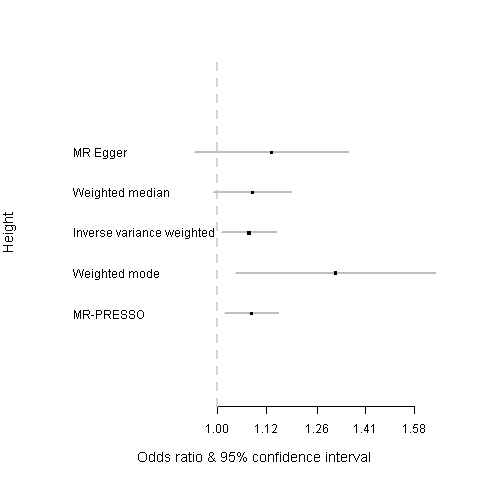 |
| 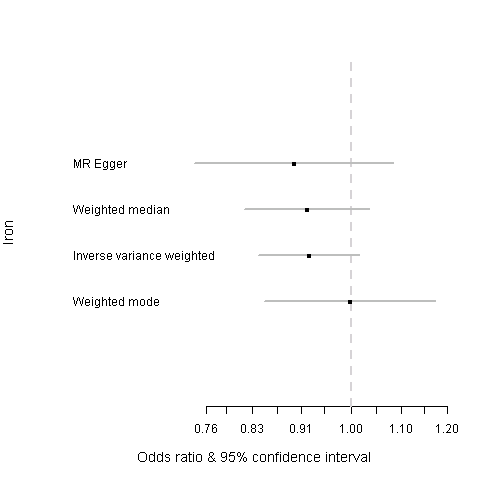 | 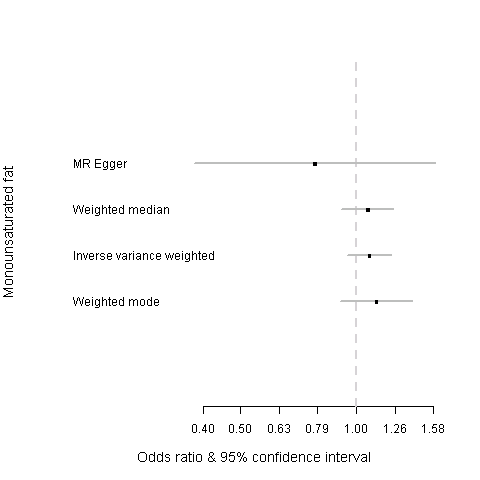 |
| 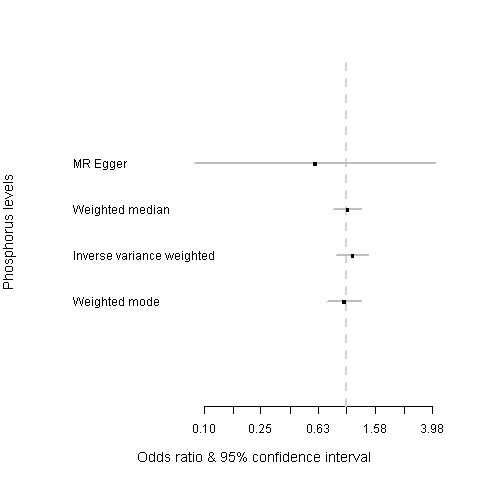 | 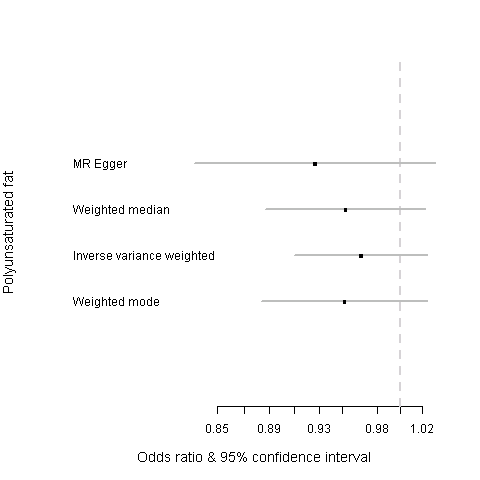 |
| 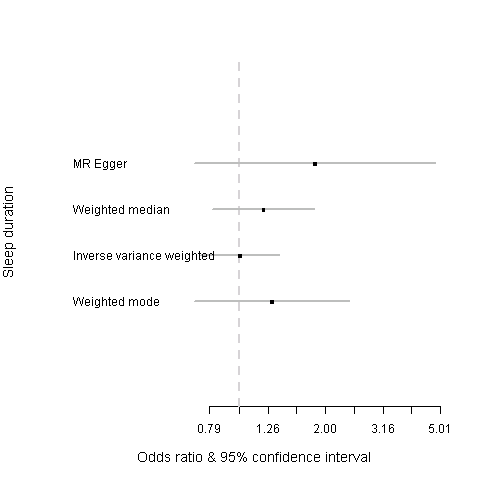 | 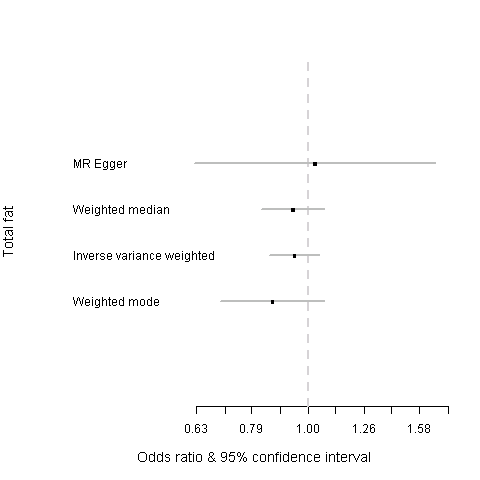 |
| 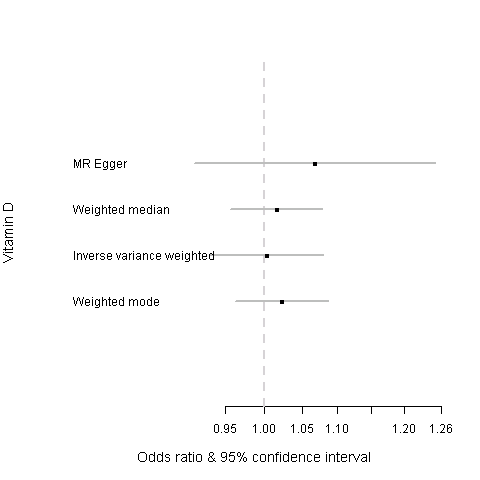 | 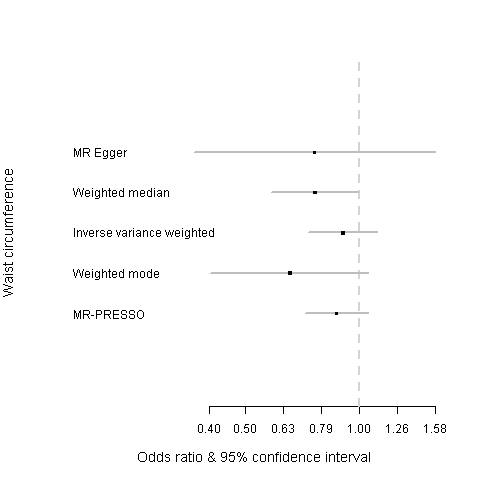 |
| 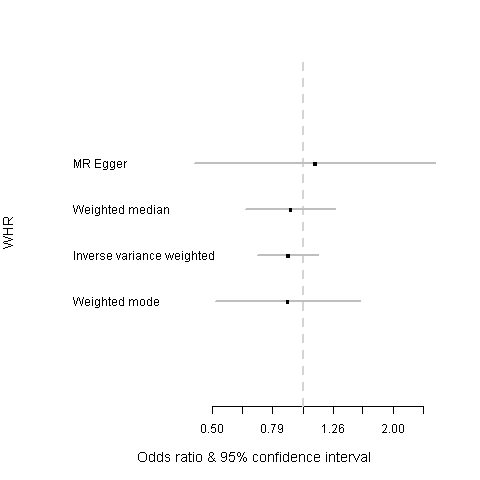 |  |

MR-PRESSO = when MR-PRESSO identified outlying SNPs for a risk factor, IVW method was re-ran after excluding these outlying SNPs.

**PIs from the PRACTICAL (**[**http://practical.icr.ac.uk/**](http://practical.icr.ac.uk/)**), CRUK, BPC3, CAPS, PEGASUS consortia:**

Rosalind A. Eeles^1,2^, Brian E. Henderson^3*^, Christopher A. Haiman^3^, ZSofia Kote-Jarai^1^, Fredrick R. Schumacher^4,5^, Sara Benlloch^6,1^, Ali Amin Al Olama^6,9^, Kenneth Muir^7,8^, Sonja I. Berndt^10^, David V. Conti^3^, Fredrik Wiklund^11^ , Stephen Chanock^10^, Susan M. Gapstur^12^, Victoria L. Stevens^12^, Catherine M. Tangen^13^, Jyotsna Batra^15,16^, Judith Clements^15,16^, Australian Prostate Cancer BioResource (APCB)^15^, Henrik Gronberg^11^ , Nora Pashayan^17,18^, Johanna Schleutker^19,20^, Demetrius Albanes^10^, Stephanie Weinstein^10^, Alicja Wolk^22, 23^, Catharine West^24^, Lorelei Mucci^25^, Géraldine Cancel-Tassin^26,27^, Stella Koutros^10^, Karina Dalsgaard Sorensen^28,29^, Eli Marie Grindedal^30^, David E. Neal^31,32^, Freddie C. Hamdy^33^, Jenny L. Donovan^34^, Ruth C. Travis^35^, Robert J. Hamilton^36^, Sue Ann Ingles^3^, Barry S. Rosenstein^37,38^, Yong-Jie Lu^39^, Graham G. Giles^40,41^, Adam S. Kibel^42^, Ana Vega^43^, Manolis Kogevinas^44,45,46,47^, Kathryn L. Penney^48^, Jong Y. Park^49^, Janet L. Stanford^50,51^, Cezary Cybulski^52^, Børge G. Nordestgaard^53,54^, Hermann Brenner^55,56,57^, Christiane Maier^58^, Jeri Kim^59^, Esther M. John^60,61^ , Manuel R. Teixeira^62,63^ , Susan L. Neuhausen^64^ , Kim De Ruyck^65^ , Azad Razack^66^ , Lisa F. Newcomb^50,67^ , Davor Lessel^68^, Radka Kaneva^69^, Nawaid Usmani^70,71^, Aswin Abraham^70,71^, Frank Claessens^72^, Paul A. Townsend^73^, Manuela Gago-Dominguez^74,75^ , Monique J. Roobol^76^, Florence Menegaux^77^, Kay-Tee Khaw^78^, Lisa Cannon-Albright^79,80^, Hardev Pandha^81^, Stephen N. Thibodeau^82^, David J Hunter^83^, Peter Kraft^83^, William J. Blot^84,85^, Elio Riboli^86.^

^*^In memorium

1	The Institute of Cancer Research, London, UK.

2	Royal Marsden NHS Foundation Trust, London, UK.

3	Department of Preventive Medicine, Keck School of Medicine, University of Southern California/Norris Comprehensive Cancer Center, Los Angeles, CA, USA.

4	Department of Population and Quantitative Health Sciences, Case Western Reserve University, Cleveland, OH, USA.

5	Seidman Cancer Center, University Hospitals, Cleveland, OH, USA.

6	Centre for Cancer Genetic Epidemiology, Department of Public Health and Primary Care, University of Cambridge, Strangeways Research Laboratory, Cambridge, UK.

7	Division of Population Health, Health Services Research and Primary Care, University of Manchester, Oxford Road, Manchester, UK.

8	Warwick Medical School, University of Warwick, Coventry, UK.

9	University of Cambridge, Department of Clinical Neurosciences, Cambridge, UK.

10	Division of Cancer Epidemiology and Genetics, National Cancer Institute, NIH, Bethesda, MD, USA.

11	Department of Medical Epidemiology and Biostatistics, Karolinska Institute, Stockholm, Sweden.

12	Epidemiology Research Program, American Cancer Society, 250 Williams Street, Atlanta, GA, USA.

13	SWOG Statistical Center, Fred Hutchinson Cancer Research Center, Seattle, WA, USA.

15	Australian Prostate Cancer Research Centre-Qld, Institute of Health and Biomedical Innovation and School of Biomedical Sciences, Queensland University of Technology, Brisbane, Queensland, Australia.

16	Translational Research Institute, Brisbane, Queensland, Australia.

17	University College London, Department of Applied Health Research, London, UK.

18	Centre for Cancer Genetic Epidemiology, Department of Oncology, University of Cambridge, Strangeways Laboratory, Cambridge, UK.

19	Institute of Biomedicine, Kiinamyllynkatu 10, FI-20014 University of Turku, Finland

20	Department of Medical Genetics, Genomics, Laboratory Division, Turku University Hospital, PO Box 52, 20521 Turku, Finland

21	Division of Cancer Epidemiology and Genetics, National Cancer Institute, NIH, Bethesda, MD, USA.

22	Division of Nutritional Epidemiology, Institute of Environmental Medicine, Karolinska Institutet, Sweden.

23	Department of Surgical Sciences, Uppsala University, Uppsala, Sweden.

24	Division of Cancer Sciences, University of Manchester, Manchester Academic Health Science Centre, Radiotherapy Related Research, The Christie Hospital NHS Foundation Trust, Manchester, M13 9PL UK.

25	Department of Epidemiology, Harvard School of Pubic Health, Boston, MA, USA.

26	CeRePP, Tenon Hospital, Paris, France

27	UPMC Sorbonne Universites, GRC N°5 ONCOTYPE-URO, Tenon Hospital, 4 rue de la Chine, Paris, France

28	Department of Molecular Medicine, Aarhus University Hospital, Denmark.

29	Department of Clinical Medicine, Aarhus University, Denmark.

30	Department of Medical Genetics, Oslo University Hospital, Norway.

31	University of Cambridge, Department of Oncology, Addenbrooke's Hospital, Cambridge, UK.

32	Cancer Research UK Cambridge Research Institute, Li Ka Shing Centre, Cambridge, UK.

33	Nuffield Department of Surgical Sciences, University of Oxford, Oxford, UK, Faculty of Medical Science, University of Oxford, John Radcliffe Hospital, Oxford, UK.

34	School of Social and Community Medicine, University of Bristol, Bristol, UK.

35	Cancer Epidemiology Unit, Nuffield Department of Population Health University of Oxford, Oxford, UK.

36	Dept. of Surgical Oncology, Princess Margaret Cancer Centre, Toronto, Canada.

37	Department of Radiation Oncology, Icahn School of Medicine at Mount Sinai, New York, NY, USA.

38	Department of Genetics and Genomic Sciences, Icahn School of Medicine at Mount Sinai, New York, NY, USA.

39	Centre for Molecular Oncology, Barts Cancer Institute, Queen Mary University of London, John Vane Science Centre, London, UK.

40	Cancer Epidemiology Division, Cancer Council Victoria, 615 St Kilda Road, Melbourne, Victoria, 3004, Australia.

41	Centre for Epidemiology and Biostatistics, Melbourne School of Population and Global Health, The University of Melbourne, Melbourne, Victoria 3010, Australia

42	Division of Urologic Surgery, Brigham and Womens Hospital, Boston, MA, USA.

43	Fundación Pública Galega de Medicina Xenómica-SERGAS, Grupo de Medicina Xenómica, CIBERER, IDIS, Santiago de Compostela, Spain.

44	ISGlobal, Barcelona, Spain

45	IMIM (Hospital del Mar Medical Research Institute), Barcelona, Spain.

46	Universitat Pompeu Fabra (UPF), Barcelona, Spain

47	CIBER Epidemiología y Salud Pública (CIBERESP), Madrid, Spain

48	Channing Division of Network Medicine, Department of Medicine, Brigham and Women's Hospital/Harvard Medical School, Boston, MA, USA.

49	Department of Cancer Epidemiology, Moffitt Cancer Center, Tampa, USA.

50	Division of Public Health Sciences, Fred Hutchinson Cancer Research Center, Seattle, Washington, USA.

51	Department of Epidemiology, School of Public Health, University of Washington, Seattle, Washington, USA.

52	International Hereditary Cancer Center, Department of Genetics and Pathology, Pomeranian Medical University, Szczecin, Poland.

53	Faculty of Health and Medical Sciences, University of Copenhagen, Denmark.

54	Department of Clinical Biochemistry, Herlev and Gentofte Hospital, Copenhagen University Hospital, Herlev, Denmark.

55	Division of Clinical Epidemiology and Aging Research, German Cancer Research Center (DKFZ), Heidelberg, Germany.

56	German Cancer Consortium (DKTK), German Cancer Research Center (DKFZ), Heidelberg, Germany.

57	Division of Preventive Oncology, German Cancer Research Center (DKFZ) and National Center for Tumor Diseases (NCT), Heidelberg, Germany.

58	Institute for Human Genetics, University Hospital Ulm, Ulm, Germany.

59	The University of Texas M. D. Anderson Cancer Center, Department of Genitourinary Medical Oncology, Houston, TX, USA.

60	Cancer Prevention Institute of California, Fremont, CA, USA.

61	Department of Health Research & Policy (Epidemiology) and Stanford Cancer Institute, Stanford University School of Medicine, Stanford, CA , USA.

62	Department of Genetics, Portuguese Oncology Institute of Porto, Porto, Portugal.

63	Biomedical Sciences Institute (ICBAS), University of Porto, Porto, Portugal.

64	Department of Population Sciences, Beckman Research Institute of the City of Hope, Duarte, CA, USA.

65	Ghent University, Faculty of Medicine and Health Sciences, Basic Medical Sciences, Gent, Belgium.

66	Department of Surgery, Faculty of Medicine, University of Malaya, Kuala Lumpur, Malaysia.

67	Department of Urology, University of Washington, Seattle, WA, USA.

68	Institute of Human Genetics, University Medical Center Hamburg-Eppendorf, Hamburg, Germany.

69	Molecular Medicine Center, Department of Medical Chemistry and Biochemistry, Medical University, Sofia, Bulgaria.

70	Department of Oncology, Cross Cancer Institute, University of Alberta, Edmonton, Alberta, Canada.

71	Division of Radiation Oncology, Cross Cancer Institute, Edmonton, Alberta, Canada.

72	Molecular Endocrinology Laboratory, Department of Cellular and Molecular Medicine, KU Leuven, Leuven, Belgium.

73	Division of Cancer Sciences, Manchester Cancer Research Centre, Faculty of Biology, Medicine and Health, Manchester Academic Health Science Centre, NIHR Manchester Biomedical Research Centre, Health Innovation Manchester, Univeristy of Manchester, UK.

74	Genomic Medicine Group, Galician Foundation of Genomic Medicine, Instituto de Investigacion Sanitaria de Santiago de Compostela (IDIS), Complejo Hospitalario Universitario de Santiago, Servicio Galego de Saúde, SERGAS, Santiago De Compostela, Spain.

75	University of California San Diego, Moores Cancer Center, La Jolla, CA, USA.

76	Department of Urology, Erasmus University Medical Center, Rotterdam, the Netherlands.

77	Cancer & Environment Group, Center for Research in Epidemiology and Population Health (CESP), INSERM, University Paris-Sud, University Paris-Saclay, Villejuif, France.

78	Clinical Gerontology Unit, University of Cambridge, Cambridge, UK.

79	Division of Genetic Epidemiology, Department of Medicine, University of Utah School of Medicine, Salt Lake City, Utah, USA.

80	George E. Wahlen Department of Veterans Affairs Medical Center, Salt Lake City, UT, USA.

81	The University of Surrey, Guildford, Surrey, UK.

82	Department of Laboratory Medicine and Pathology, Mayo Clinic, Rochester, MN, USA.

83	Program in Genetic Epidemiology and Statistical Genetics, Department of Epidemiology, Harvard T.H. Chan School of Public Health, Boston, MA, USA.

84	Division of Epidemiology, Department of Medicine, Vanderbilt University Medical Center, TN, USA.

85	International Epidemiology Institute, Rockville, MD, USA

86	Department of Epidemiology and Biostatistics, School of Public Health, Imperial College London, SW7 2AZ, UK

CRUK and PRACTICAL consortium

This work was supported by the Canadian Institutes of Health Research, European Commission's Seventh Framework Programme grant agreement n° 223175 (HEALTH-F2-2009-223175), Cancer Research UK Grants C5047/A7357, C1287/A10118, C1287/A16563, C5047/A3354, C5047/A10692, C16913/A6135, and The National Institute of Health (NIH) Cancer Post-Cancer GWAS initiative grant: No. 1 U19 CA 148537-01 (the GAME-ON initiative). This work was awarded by Prostate Cancer Canada and is proudly funded by the Movember Foundation – Grant # D2013-36.

We would also like to thank the following for funding support: The Institute of Cancer Research and The Everyman Campaign, The Prostate Cancer Research Foundation, Prostate Research Campaign UK (now Prostate Action), The Orchid Cancer Appeal, The National Cancer Research Network UK, The National Cancer Research Institute (NCRI) UK. We are grateful for support of NIHR funding to the NIHR Biomedical Research Centre at The Institute of Cancer Research and The Royal Marsden NHS Foundation Trust.

The Prostate Cancer Program of Cancer Council Victoria also acknowledge grant support from The National Health and Medical Research Council, Australia (126402, 209057, 251533, , 396414, 450104, 504700, 504702, 504715, 623204, 940394, 614296,), VicHealth, Cancer Council Victoria, The Prostate Cancer Foundation of Australia, The Whitten Foundation, PricewaterhouseCoopers, and Tattersall’s. EAO, DMK, and EMK acknowledge the Intramural Program of the National Human Genome Research Institute for their support.

Genotyping of the OncoArray was funded by the US National Institutes of Health (NIH) [U19 CA 148537 for ELucidating Loci Involved in Prostate cancer SuscEptibility (ELLIPSE) project and X01HG007492 to the Center for Inherited Disease Research (CIDR) under contract number HHSN268201200008I]. Additional analytic support was provided by NIH NCI U01 CA188392 (PI: Schumacher).

Funding for the iCOGS infrastructure came from: the European Community's Seventh Framework Programme under grant agreement n° 223175 (HEALTH-F2-2009-223175) (COGS), Cancer Research UK (C1287/A10118, C1287/A 10710, C12292/A11174, C1281/A12014, C5047/A8384, C5047/A15007, C5047/A10692, C8197/A16565), the National Institutes of Health (CA128978) and Post-Cancer GWAS initiative (1U19 CA148537, 1U19 CA148065 and 1U19 CA148112 - the GAME-ON initiative), the Department of Defence (W81XWH-10-1-0341), the Canadian Institutes of Health Research (CIHR) for the CIHR Team in Familial Risks of Breast Cancer, Komen Foundation for the Cure, the Breast Cancer Research Foundation, and the Ovarian Cancer Research Fund.

BPC3

The BPC3 was supported by the U.S. National Institutes of Health, National Cancer Institute (cooperative agreements U01-CA98233 to D.J.H., U01-CA98710 to S.M.G., U01-CA98216 toE.R., and U01-CA98758 to B.E.H., and Intramural Research Program of NIH/National Cancer Institute, Division of Cancer Epidemiology and Genetics).

CAPS

CAPS GWAS study was supported by the Cancer Risk Prediction Center (CRisP; www.crispcenter.org), a Linneus Centre (Contract ID 70867902) financed by the Swedish Research Council, (grant no K2010-70X-20430-04-3), the Swedish Cancer Foundation (grant no 09-0677), the Hedlund Foundation, the Soederberg Foundation, the Enqvist Foundation, ALF funds from the Stockholm County Council. Stiftelsen Johanna Hagstrand och Sigfrid Linner's Minne, Karlsson's Fund for urological and surgical research.

PEGASUS

PEGASUS was supported by the Intramural Research Program, Division of Cancer Epidemiology and Genetics, National Cancer Institute, National Institutes of Health.

22.	**Diet, nutrition, physical activity and prostate cancer.** *World Cancer Research fund/American Institute For Cancer Research Continuous Update Project Expert Report 2018*.

23.	World Cancer Research Fund/ American Institute of Cancer Research. Food, Nutrition, Physical Activity, and the Prevention of Cancer: a Global Perspective. Washington DC: AICR, 2007 (Second Expert Report). 2007.
